# Microtubule-associated protein 1S (MAP1S): a cardioprotective factor against post-myocardial infarction remodeling via apoptosis inhibition

**DOI:** 10.64898/2026.09.16.752033

**Authors:** Pia Morales, Yulia S Kohar, Efta Triastuti, Ardiansah Bayu Nugroho, Min Zi, Sukhpal Prehar, Danielle Purewal, Gina Galli, Leyuan Liu, Elizabeth J Cartwright, Delvac Oceandy

## Abstract

Adverse cardiac remodeling following myocardial infarction (MI) are driven by processes including autophagy and apoptosis. While microtubule-associated protein 1S (MAP1S) is known to regulate autophagy, its role in the cardiac pathological conditions remains unclear. This study aimed to elucidate the role of MAP1S and its underlying mechanisms in pathological cardiac remodeling. Following MI, MAP1S knockout mice exhibited increased mortality, impaired cardiac function, and elevated apoptosis. Similarly, MAP1S silencing in cultured cardiomyocytes augmented apoptosis under oxidative stress. Mechanistic investigation revealed potential link to the Hippo pathway. MAP1S knockdown in cardiomyocytes increased Mammalian Ste-20 like 1/2 (MST1/2) activation and reduced Yes-associated protein (YAP) activity, potentially explaining apoptosis regulation. Conversely, MAP1S overexpression reduced apoptosis and positively modulated autophagy. Importantly, in vivo modRNA-mediated MAP1S overexpression protected against apoptosis and adverse remodeling. This study reveals that MAP1S protects the heart from excessive apoptosis and adverse remodeling following MI, likely by modulating the Hippo signaling pathway.

## INTRODUCTION

Ischaemic heart disease (IHD) is one of the main causes of mortality and morbidity worldwide, and it is estimated that the global burden due to IHD continues to increase by 2050^1^. Heart attack, or acute myocardial infarction (MI), which is the acute manifestation of IHD, is one of the most common causes of heart failure (HF)^2^. A recent study showed that around 20-25% of patients with acute MI developed HF within 1 year^3^, indicating a very high risk of HF development among MI survivors.

In response to ischaemic injury, complex molecular, cellular, and extracellular matrix changes occur in the heart, which together alter the morphology and function of the heart. This process, called cardiac remodeling, is critical in determining the clinical outcomes caused by the primary cardiac injuries such as MI. This is because progressive adverse remodeling can lead to the enlargement of the heart, dilatation of the cardiac chambers, and significant reduction in contractile function, which eventually results in heart failure^4^. Thus, a complete understanding of the molecular and cellular mechanisms underlying cardiac remodeling is very important in order to find new strategies to control the progression of this adverse process and prevent the development of heart failure.

It has been widely known that autophagy is involved in mediating cardiac remodeling. Autophagy is a mechanism to degrade and recycle damaged cellular materials, such as misfolded proteins and dysfunctional organelles (reviewed in^5^). Clearance of damaged organelles and misfolded proteins is of particular importance during cardiac injury because accumulation of damaged organelles (e.g. mitochondria) may trigger apoptosis, necrosis, and oxidative stress^6^.

Several studies have characterized the roles of key autophagy regulators in cardiac pathological conditions. Most of the studies have investigated the importance of molecules that are involved in autophagy initiation and autophagic vesicle formation during the progression of pathological heart remodeling. For example, the roles of Atg5, Beclin1, and Mst1, which are important factors in the regulation of autophagy initiation, have been characterized in the setting of cardiac pathological conditions, such as myocardial infarction and aortic constriction^7, 8, 9, 10^. These observations have indicated that reduction of autophagic flux during cardiac pathological conditions was detrimental.

Despite ample observations on the role of molecules that are involved in autophagy initiation, there is only limited knowledge on the regulation of cargo recognition and binding to autophagic vacuoles. Recently, several molecules that are capable of linking damaged organelles to the autophagosomes have been identified, one of which is the microtubule-associated protein 1S (MAP1S)^11^. MAP1S is a member of the microtubule-associated proteins (MAP) family, which are conventionally regarded as components of the cytoskeletal networks. Most members of this protein family mediate interactions between microtubules and actin, stabilize microtubules, and are involved in regulating microtubule dynamics^12^. In addition to these functions, isoform 1S of this protein (MAP1S) may have additional roles in regulating autophagy. MAP1S interacts with one of the main regulators of autophagy, the microtubule-associated protein light chain 3 (LC3)^11^, which, if activated, is cleaved and conjugated with phosphatidylethanolamine to form membrane-associated LC3-II. On the other hand, MAP1S also binds to the leucine-rich pentatricopeptide repeat-containing protein (LRPPRC)^13^, which is known as a mitochondrion-associated protein. Based on this evidence, it has been postulated that MAP1S might play an essential role in mitophagy by bridging autophagy-related proteins and autophagic membranes to the damaged mitochondria; however, its role in the heart, in particular in pathological conditions, is not completely understood. Therefore, in this study we focused on investigating the role of MAP1S in mediating cardiac remodeling during ischaemic cardiac injury, which is the most common cause of cardiovascular morbidity and mortality.

## RESULTS

### MAP1S expression in the heart

To understand MAP1S’ functions in the heart, we first examine its expression in normal and pathologic conditions. We detected MAP1S expression in both cardiomyocytes and cardiac fibroblasts isolated from neonatal rat hearts. The 126 kDa full-length and the active form heavy chain fragment of MAP1S (∼100 kDa) were detected by Western blots in both types of cardiac cells (Fig. 1A). MAP1S expression in adult mouse heart was evident and in a mouse model of *Map1s* ablation^11^ the cardiac expression of this protein was markedly depleted (Fig. 1B).

**Figure 1.**
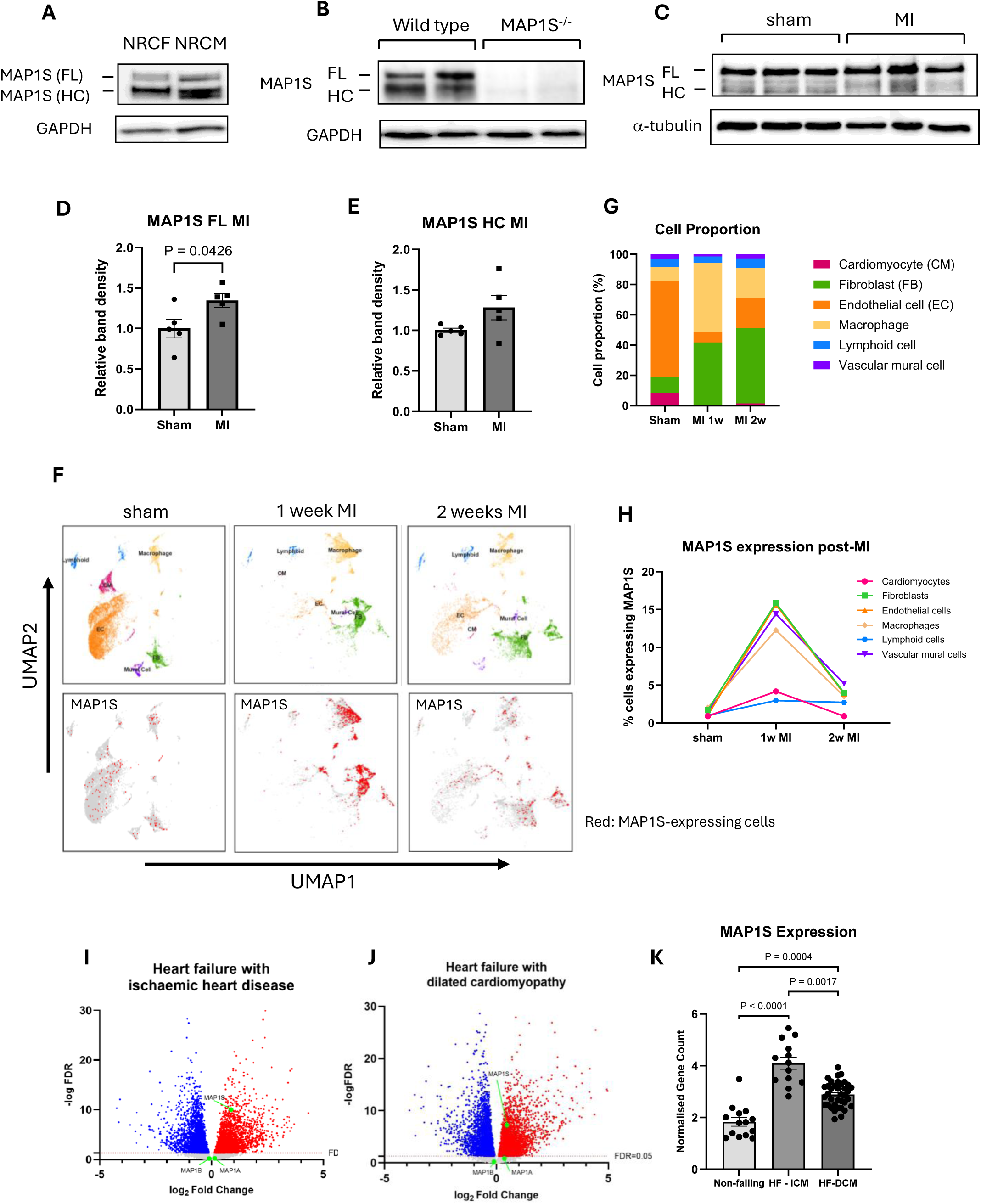
MAP1S cardiac expression in normal and pathological conditions. **A)** Representative Western blots showing expressions of MAP1S full-length protein (MAP1S-FL) and heavy chain (MAP1S-HC) fragment in neonatal rat cardiomyocytes (NRCM) and cardiac fibroblasts (NRCF)(n = 3 independent cell preparations). **B)** Western blot analysis demonstrated that both MAP1S-FL and MAP1S-HC are expressed in adult wild-type mouse hearts and the expression is completely ablated in MAP1S global knockout (MAP1S^-/-^) hearts. **C)** Representative Western blots and **(D, E)** quantification of band density revealed that in response to myocardial infarction (MI), MAP1S-FL was significantly increased; however, no significant difference was observed in MAP1S-HC. (n = 5 in each group). **F)** Analysis of single cell RNAseq data of mouse hearts following MI (GEO database assession no: GSE163956). UMAP plots indicate clustering of cardiac cells and expression of MAP1S comparing sham, 1 week MI and 2 weeks MI. **G)** Proportion of each cell types in the heart of sham, 1 week MI, and 2 weeks MI mice. **H)** Expression of MAP1S in each cell types following MI. **I-J)** Volcano plots of differentially expressed genes in human failing hearts with **I)** ischaemic heart disease (HF-ICM) and **J)** dilated cardiomyopathy (HF-DCM). Green dots represent expressions of MAP1 family proteins (MAP1A, MAP1B and MAP1S). **K**) Analysis of normalised gene counts showed significant increase in cardiac MAP1S expression in patients with HF-ICM and HF-DCM)(HF-ICM, n=13; HF-DCM, n=37; non-failing, n=14; data analysed from GEO database assession no:GSE116250). Statistical tests used: (**D,E**) Student’s *t-test,* (**K**) Kruskall-Wallis test followed by multiple comparisons test. Numbers above the bars indicate *P* values.

We then assessed cardiac expression of MAP1S in wild type (WT) mice following pathological stress. Following myocardial infarction (MI) for 4 weeks, we found a significant increase of MAP1S full length expression in MI group, but not the heavy chain (Fig. 1C-E).

To examine cell specific expression of MAP1S during MI, we analysed single cell transcriptomic dataset in the GEO database (GSE163956). Our analysis revealed changes in cell composition following MI in mice (Fig.1F-G). MAP1S is expressed in most of cardiac cells including fibroblasts, macrophages, cardiomyocytes and endothelial cells. Interestingly, the proportion of cells expressing MAP1S was markedly elevated at 1 week post-MI; however, its expression was reduced to those comparable to basal condition at 2 weeks post-MI (Fig.1F,H).

The data above suggest that MAP1S may play important role in animal model of MI. However, it is important to investigate expressions in human hearts in pathological conditions. Therefore, we analysed GEO datasets GSE116250, which consists of gene expression datasets of human left ventricular tissues from heart failure participants with ischaemic cardiomyopathy, dilated cardiomyopathy and non-failing heart as controls. Analysis of volcano plots was performed to evaluate expressions of MAP1 family of proteins in these conditions. As shown in figures 1I-J MAP1S expression was markedly elevated in HF participants with ICM and DCM. However, MAP1A and MAP1B expressions were less significantly altered in these conditions. Analysis of normalised gene count revealed a significant increase in MAP1S expression in HF individuals with ICM and DCM (Fig. 1K). Overall, our data demonstrate that MAP1S expression is induced in cardiac pathological settings in both mouse models and in human heart diseases, which might suggest its essential roles during these conditions.

### MAP1S ablation reduced mouse survival following myocardial infarction

To determine the effect of MAP1S ablation during ischaemic injury, we subjected MAP1S^-/-^ mice and WT littermates to MI. We observed a significant reduction of mouse survival in the MAP1S^-/-^ group within the first week of MI (MAP1S^-/-^, 46% survival vs WT, 70% survival, Fig. 2A) although the extent of cardiac injury due to coronary artery ligation was comparable between these groups as indicated by the serum cTnT level at 24 hours post ligation (Fig. 2B). In keeping with this data, the extent of infarct size in surviving MAP1S knockout animals at 4 weeks post-MI did not differ compared to those of WT mice (Fig. 2C-D).

**Figure 2.**
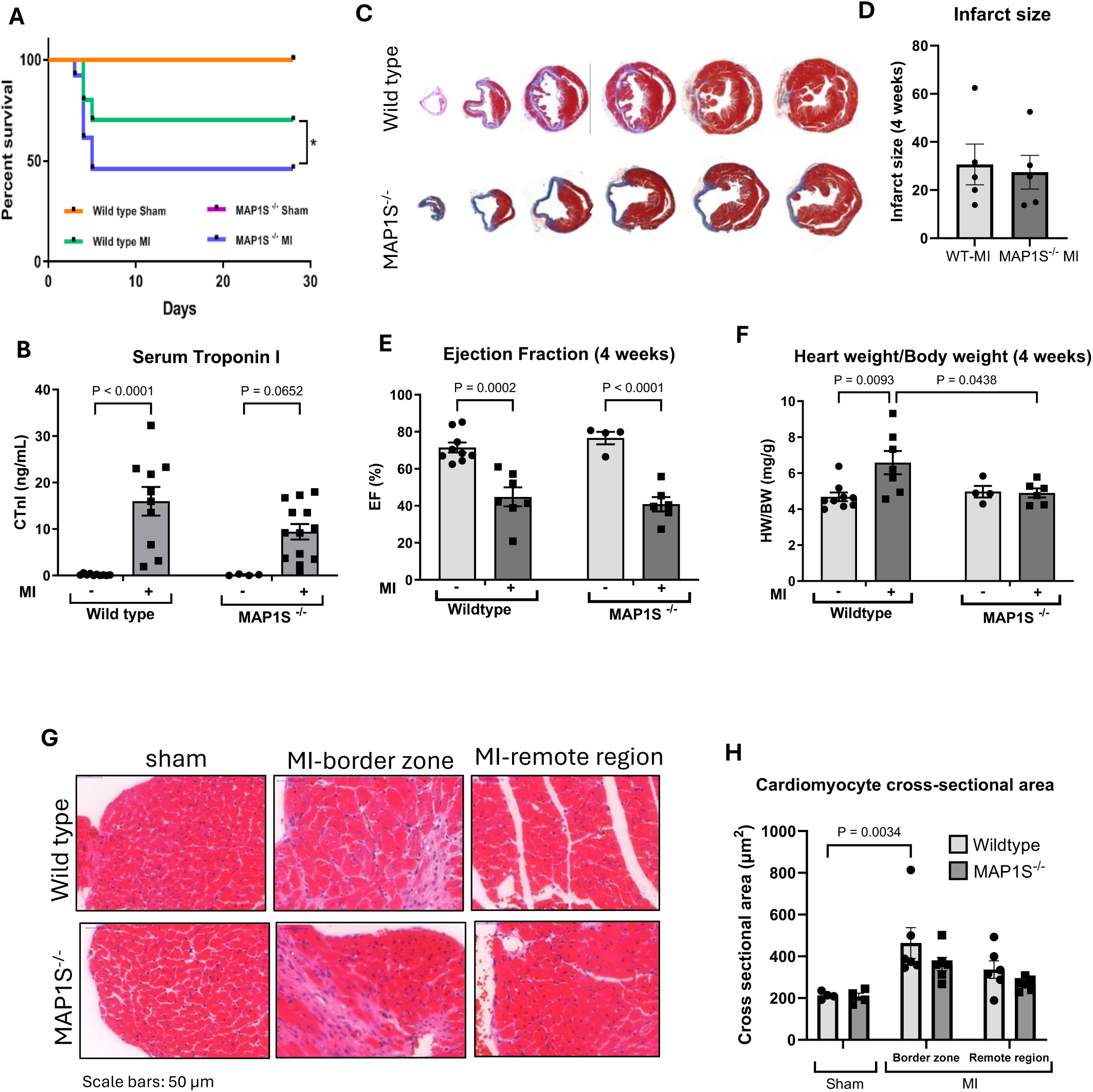
MAP1S deletion increased mortality and apoptosis in mice with myocardial infarction. MAP1S^-/-^ mice and wild-type (WT) littermates were subjected to myocardial infarction (MI) for 4 weeks. **A)** MAP1S^-/-^ mice exhibited reduced survival within the first week of MI. (WT sham, n = 9 for WT MI, n = 7 for MAP1S^-/-^ sham, n = 4 for MAP1S^-/-^ MI, n=6). **B)** Serum cardiac troponin I levels at 24 hours post-MI. **C)** Representative Masson’s trichrome-stained cardiac sections from WT and MAP1S^-/-^ hearts and **D)** quantification of infarct size showed no difference between the two MI groups at 4 weeks after MI. **E)** Analysis of ejection fraction revealed no difference between WT and MAP1S^-/-^ mice at 4 weeks post-MI. However, **F)** analysis of heart weight/body weight ratio indicated reduced heart size in MAP1S^-/-^ mice at 4 weeks following MI. **G)** Haematoxylin and eosin-stained sections and **H)** quantification of cardiomyocyte cross-sectional area showed a significant increase in WT mice at the infarct border zone, but not in MAP1S^-/-^ mice. There was no change in cardiomyocyte size in the remote zone. Data are presented as mean ± SEM. Statistical tests used: (**B,D,E,F,H**) Two-way ANOVA followed by posthoc multiple comparisons. Numbers above the bars indicate *P* values.

To further assess the phenotype of surviving mice at 4 weeks post-MI, we conducted echocardiography analysis. Despite a significant reduction in cardiac contractility between the sham and MI groups, there was no significant difference in ejection fraction (EF) between MAP1S^-/-^ and WT mice both basally and at 4 weeks after MI (Fig. 2E). Assessment of other echocardiography parameters such as chamber dimension and wall thickness did not show any difference between knockout and WT mice (Supplementary Fig.1). However, we observed a significantly smaller heart weight/body weight (HW/BW) ratio in MAP1S^-/-^ mice compared to WT at 4 weeks after MI (Fig. 2F). Consistently, an analysis of the cardiomyocyte cross-sectional area of histological sections suggested a trend of smaller cardiomyocyte size in the infarct border zone but not in the remote zone of MAP1S^-/-^ MI mice compared with the WT-MI group (Fig. 2G-H).

### Cardiac apoptosis level was elevated in MAP1S^-/-^ mice following MI

To examine the level of myocardial apoptotic cell death, we performed TUNEL analysis on cardiac sections at 4 weeks post-MI. As indicated in figure 3A-B, apoptosis was more prevalent in MAP1S^-/-^ hearts compared to WT following MI. Thus, it is possible that the cause of excessive mortality in MAP1S^-/-^ mice during the first week of MI was due to enhanced apoptotic cell death. To test this hypothesis, we performed additional MI experiments to analyse apoptosis level in the heart at day 3 post-MI, since the majority of death occurred at day 3-5 post-MI. Strikingly, the number of apoptotic cardiomyocytes was markedly higher in MAP1S^-/-^ mice at day 3 post-MI (Fig.3C-D). Overall, these data suggest that excessive apoptosis might contribute to the higher mortality rate in MAP1S^-/-^ mice after MI.

**Figure 3.**
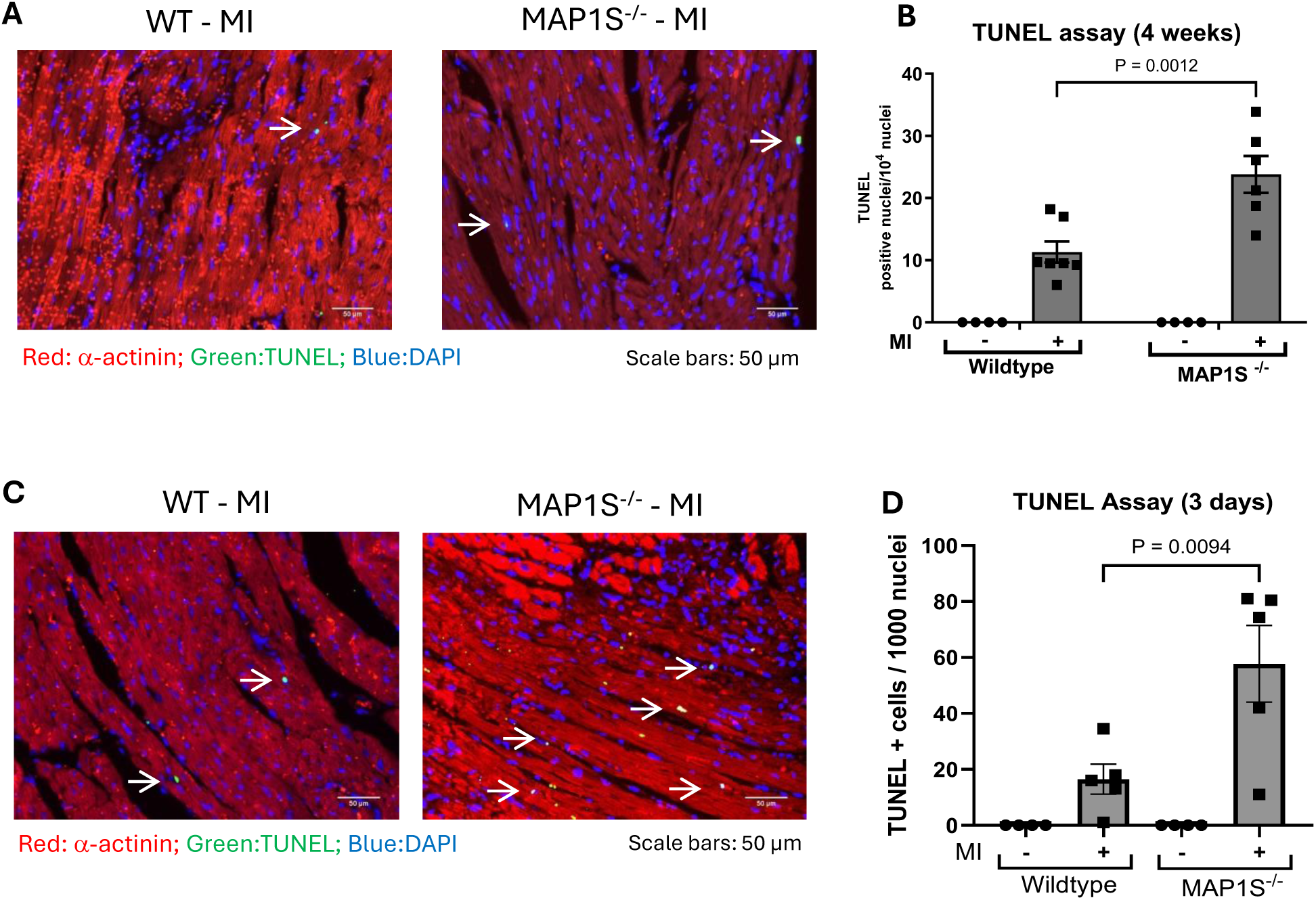
MAP1S deletion increases apoptosis during myocardial infarction. **A)** Representative images of TUNEL-stained cardiac tissue sections of MAP1S-/-and WT mice following MI for 4 weeks. **B)** Quantification of TUNEL-positive cells demonstrated enhanced apoptosis in MAP1S^-/-^ hearts at 4 weeks after MI. (WT sham, n = 4; WT MI, n=7, MAP1S^-/-^ sham, n = 4; MAP1S^-/-^ MI, n=6). We then performed TUNEL analysis on heart sections of mice at 3 days after MI. **C)** Representative TUNEL staining and **D)** quantification of TUNEL-positive cells demonstrated significantly increased apoptosis in MAP1S^-/-^ hearts 3 days after MI. (WT sham, n=4; WT MI,n=5; MAP1S^-/-^ sham,n=4; MAP1S^-/-^ MI, n=5). Statistical tests used: (**B,D**) Two-way ANOVA followed by posthoc multiple comparisons. Numbers above the bars indicate *P* values.

### Effects of MAP1S gene silencing on autophagy and apoptosis regulation in cardiomyocytes

To further study the mechanism of MAP1S-mediated apoptosis in cardiomyocytes, we created a cellular model of MAP1S deficiency using siRNA-mediated gene silencing in cultured neonatal rat cardiomyocytes (NRCM). As shown in figure 4A, transfection with siRNA targeting MAP1S resulted in a marked reduction in MAP1S expression in NRCM.

**Figure 4.**
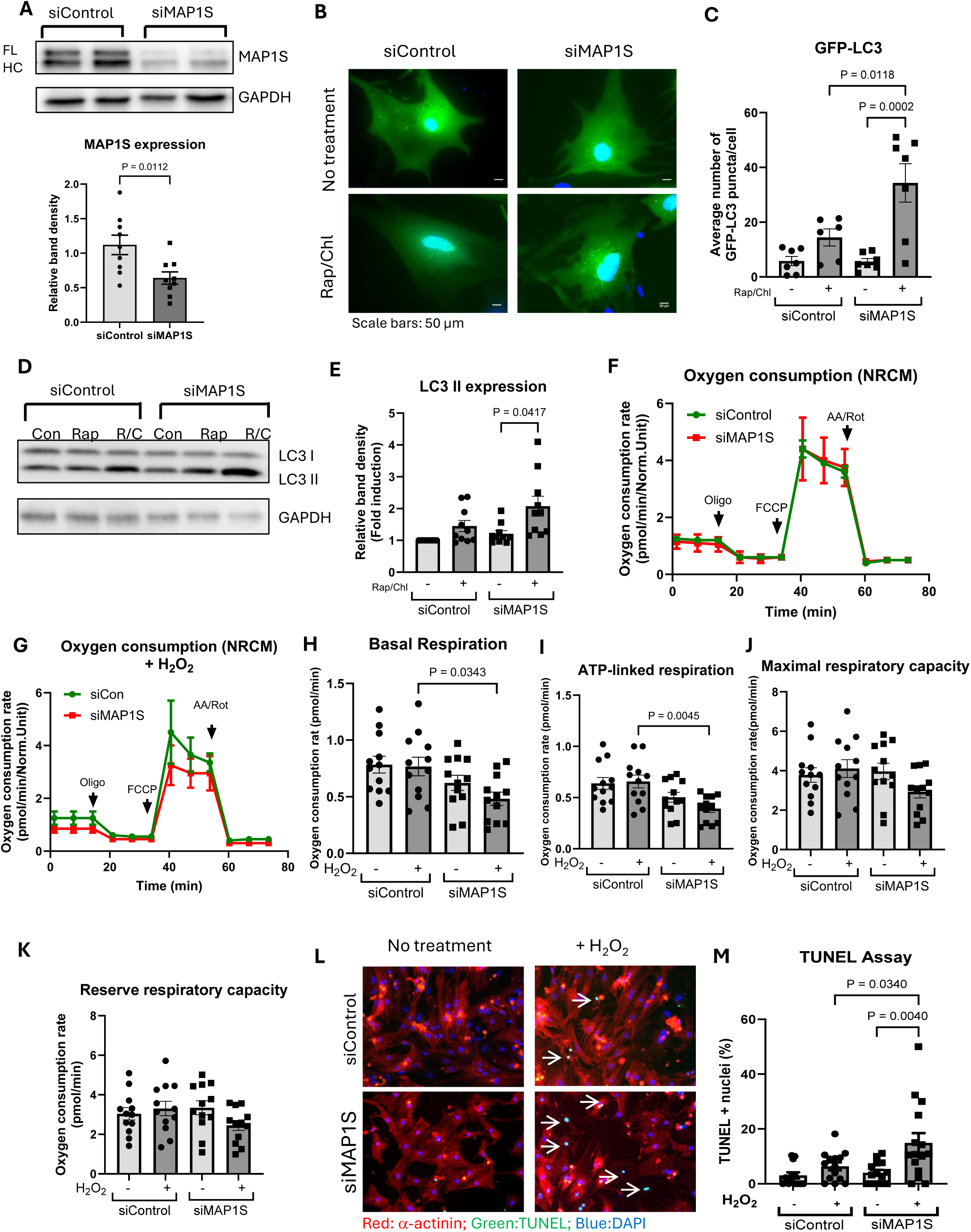
MAP1S gene silencing affects autophagy and apoptosis in NRCM. **A)** Western blots and band density quantification showing reduced MAP1S expression in NRCM following transfection with MAP1S-targeting siRNA (siMAP1S)(n = 9 independent experiments). **B)** Images of MAP1S-depleted NRCM expressing GFP-tagged LC3 and co-treated with rapamycin (5 μM) and chloroquine (3 μM) for 2 hours. **C)** Quantification of GFP-LC3 puncta per cell (n=7 independent experiments). **D** Western blots and **E)** band density quantification of endogenous LC3-II in MAP1S-depleted NRCMs after rapamycin and chloroquine co-treatment (n=10 independent experiments). **F)** Representative OCR traces of Seahorse analysis of MAP1S-depleted vs control NRCM under basal conditions and **G)** in the presence of 25 μM H_2_O_2_. Analysis of **H)** basal respiration and **I)** ATP synthesis-linked respiration indicated that NRCM lacking MAP1S displayed significantly lower OCR in the presence of H_2_O_2_. While no difference in OCR was observed during **J**) maximal respiration and **K)** reserve respiration. (n = 3 independent experiments; Con, control; Rap, rapamycin; R/C, rapamycin and chloroquine; Oligo, oligomycin; FCCP, carbonyl cyanide-4-(trifluoromethoxy) phenylhydrazone; AA: antimycin A; Rot, rotenone). **L)** TUNEL staining and **M)** quantification of TUNEL-positive cells showed increased apoptosis in MAP1S-depleted NRCMs following H_2_O_2_ treatment (100 μM, 2 hours) (n = 13-15 independent experiments). Statistical tests used: (**A**) Student’s *t-test*; (**C,H,I,J,K,M**) One-way ANOVA followed by posthoc multiple comparisons. (**E**) Kruskal-Wallis test followed by posthoc multiple comparisons. Numbers above the bars indicate *P* values.

Previous reports have associated MAP1S with the regulation of autophagy^11, 14^, and studies indicated that ablation of this protein led to the accumulation of autophagosomes due to the reduction in binding to lysosomes. We therefore used the GFP-LC3 construct to test if MAP1S ablation in NRCM affects autophagic flux. Data shown in figures 4B-C indicated an increased number of GFP-LC3 puncta following rapamycin/chloroquine treatment in MAP1S-deficient myocytes, suggesting the accumulation of autophagosomes, a result that is consistent with previous report^11^. Western blot analysis of LC3II level supported this finding, as we observed a trend of higher LC3II level in NRCM-lacking MAP1S (Fig. 4D–E).

Interestingly, analysis of electron micrographs of heart tissues indicated a trend of reduced autolysosome/autophagosome ratio in MAP1S^-/-^ mice after MI (Supplementary Fig. 2A-B), suggesting a possible alteration in autophagosome-lysosome fusion. This is consistent with the in vitro data above showing the accumulation of autophagic puncta in MAP1S-deficient NRCM.

We then analysed the effects of MAP1S gene silencing on mitochondrial functions since mitochondria is known as a substrate of autophagy. Analysis of mitochondrial oxygen consumption rate in cultured neonatal cardiomyocytes using the Seahorse system suggested similar levels of oxygen consumption rates (OCRs) at basal conditions between MAP1S-deficient NRCM compared to control cells. However, following induction of oxidative stress using H2O2 we observed a reduction of mitochondrial respiratory capacity in MAP1S-depleted NRCM (Fig.4F-K and Supplementary Fig.2C).

Importantly, in consistent with the phenotypes of MAP1S^-/-^ mice in response to MI, TUNEL analysis showed that deficiency of MAP1S in cultured cardiomyocytes led to the induction of apoptosis in response to oxidative stress due to treatment with H2O2 (Fig.4L-M). Analysis of the expressions of apoptosis markers demonstrated a trend of higher caspase-3 and cytochrome-c. However, the level of the mitochondria-related apoptosis mediators (Bax, Bad, Bcl-2, Bcl-xL) in cardiomyocytes did not seem to be affected by MAP1S gene silencing (Supplementary Fig.3), which is consistent with the in vivo data. These suggest that there might be autophagy-independent mechanism for apoptosis regulation by MAP1S in cardiomyocytes.

### MAP1S is associated with the Hippo signaling pathway

To investigate the mechanism underlying the phenotypes of MAP1S-deficient mice and cardiomyocytes, we performed in silico analysis of known protein interaction partners of MAP1S (Supplementary Fig. 4A). We identified 48 experimentally validated MAP1S interacting proteins (Supplementary Table 2). The network of interacting proteins is shown in Supplementary figure 6B. Of the 48 MAP1S interacting partners, 46 of them are expressed in the heart according to the Expression Atlas^15^. Based on the number of supporting evidence, 13 out of 46 MAP1S interacting proteins have been confirmed by more than one experimental evidence (Supplementary Table 2).

We then performed annotation enrichment analysis of these 13 proteins using the KEGG and Reactome databases to identify signaling pathways that are over-represented in the list of MAP1S interacting partners. As shown in Supplementary figures 4C-D, the Hippo pathway appeared to be the most significantly over-represented signaling pathway with the strongest P-value and highest number of interacting partners, indicating that this pathway might be associated with and regulated by MAP1S.

### MAP1S gene silencing in NRCM altered the Hippo pathway and YAP activation

The finding above prompted us to evaluate expressions and activities of key components of the Hippo pathway in the setting of MAP1S gene deletion. We first analysed the level of phosphorylated MST1/2, which is the key core component of the Hippo pathway. We found that depletion of MAP1S increased phosphorylation of MST1/2 in NRCM, indicating increased activity of the Hippo pathway (Fig. 5A-B). The main downstream effector of this pathway is YAP, which is negatively modulated by the active Hippo pathway. Indeed, analysis of active YAP levels and YAP nuclear translocation suggested a reduction of YAP activity following MAP1S knockdown (Fig. 5C-F). Moreover, YAP transcriptional activity was also reduced, as shown by the YAP-luciferase assay (Fig. 5G).

**Figure 5.**
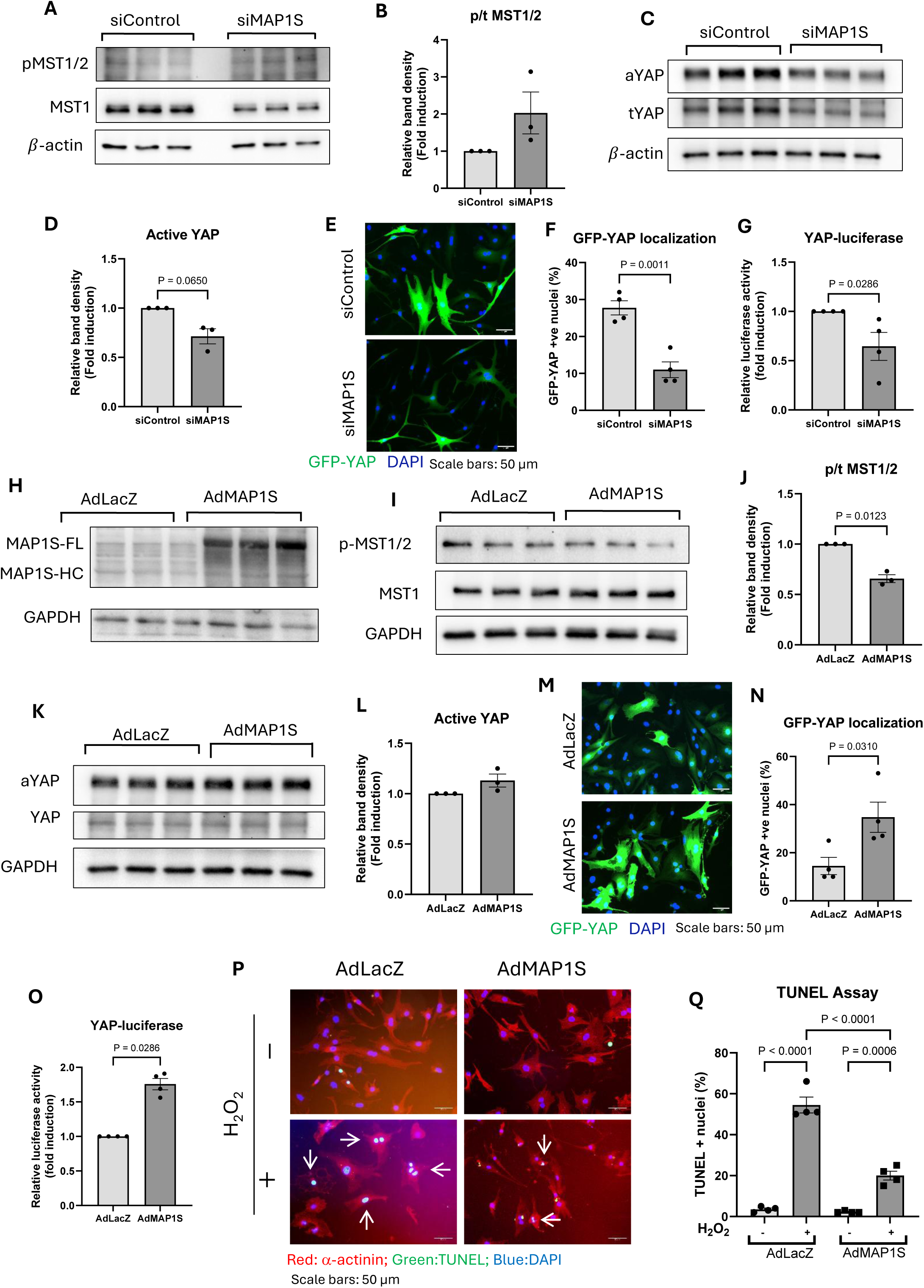
MAP1S modulates Hippo pathway in cardiomyocytes. **A)** Western blots and **B)** quantification of band density of phosphorylated/total MST1/2 in MAP1S-deficient NRCMs indicated increased MST1/2 phosphorylation in NRCM lacking MAP1S (n=3 independent experiments). **C)** Western blots and **D)** band density quantification of active/total YAP showed reduced levels of active YAP in MAP1S-deficient NRCMs (n=3 independent experiments). **E)** YAP nuclear translocation was monitored using GFP-YAP construct. **F)** Quantification of nuclear GFP-YAP-positive cells showed a significant reduction of nuclear YAP localization in MAP1S-deficient NRCM (n=4 independent experiments). **G)** YAP-luciferase assay indicated reduced YAP transcriptional activity in MAP1S-deficient NRCMs (n=4 independent experiments). **H**) Western blots showing enhanced MAP1S expression in NRCMs following transduction with adenovirus expressing MAP1S (AdMAP1S). **I)** Western blot and **J)** quantification of band density revealed that MAP1S overexpression significantly reduced phosphorylated MST1/2 levels in NRCM (n=3 independent experiments). **K)** Western blot and **L)** band density analysis of active/total YAP showed that MAP1S overexpression did not affect the expression of active YAP in NRCM (n=3 independent experiments). However, **M)** analysis of GFP-YAP localisation and **N)** quantification of nuclear GFP-YAP-positive cells showed a significant increase in nuclear YAP localisation in MAP1S-overexpressing NRCMs (n = 4 independent experiments). **O)** YAP-luciferase assay confirmed increased YAP transcriptional activity in MAP1S-overexpressing cells (n=4 independent experiments). **P)** TUNEL staining and **Q)** quantification of TUNEL-positive cells showed significantly reduced apoptosis in MAP1S-overexpressing NRCMs following H_2_O_2_ treatment (100 μM, 2 hours) (n=4 independent experiments). Statistical tests used: (**B, D, J, L**) One sample *t-test*; (**F, N**) Student’s t-test; (**G,O**) Mann Whitney U test; (**Q**) One-way ANOVA followed by posthoc multiple comparisons. Numbers above the bars indicate *P* values.

Overall, our data suggested that the Hippo/YAP pathway might be modulated by MAP1S. Since the Hippo pathway and YAP are strong regulators of apoptosis, modulation of the Hippo/YAP pathway might explain the pro-apoptotic phenotype of MAP1S deletion.

### MAP1S overexpression in NRCM induces YAP activation

Based on the observations using MAP1S knockout mouse and siRNA-mediated gene silencing models, the logical hypothesis would be that increasing MAP1S expression in cardiomyocytes would modulate Hippo/YAP pathway and would be beneficial in reducing apoptosis and increasing cell survival following stress. To test this idea, we generated an adenovirus carrying MAP1S cDNA. Treatment with this virus successfully induced the expressions of MAP1S in NRCM (Fig. 5H).

We then examined MST1/2 phosphorylation and YAP activation in these cells. Consistent with the data from gene silencing experiments, MAP1S overexpression reduced MST1/2 phosphorylation and increased active YAP levels (Fig. 5I-L). In keeping with these results, YAP nuclear translocation and YAP-luciferase activity were also significantly elevated in NRCM overexpressing MAP1S compared to controls (Fig. 5M-O).

### MAP1S overexpression inhibits apoptosis in NRCM

Inhibition of MST1/2 and activation of YAP might be protective against apoptotic cell death. We therefore conducted TUNEL assays to test the effects of MAP1S overexpression on NRCM apoptosis following oxidative stress. As shown in figure 5P-Q, MAP1S expression was protective in NRCM treated with H2O2 since there was a significantly lower number of apoptotic cells in this experimental group. However, consistent with results from the gene silencing experiments we did not find any difference in the expressions of several mitochondria-related intrinsic mediators of apoptosis, such as Bax, Bad, Bcl-2, and Bcl-xL (Supplementary Fig. 5).

### Overexpression of MAP1S increases autophagy in NRCM

Since MAP1S is tightly associated with autophagy regulation, we next assessed the autophagy process in NRCM overexpressing MAP1S. Assessment of LC3 II using Western blot showed that MAP1S overexpression increased LC3 II expression, and more importantly, the delta LC3 II (the change in LC3 II expression after rapamycin/chloroquine stimulation) was significantly higher in NRCM overexpressing MAP1S, indicating an enhanced autophagic flux (Fig.6A-C). Analysis using the GFP-LC3 construct supported this observation since NRCM overexpressing MAP1S displayed more GFP-LC3 puncta after rapamycin/chloroquine treatment (Fig. 6D-F).

**Figure 6.**
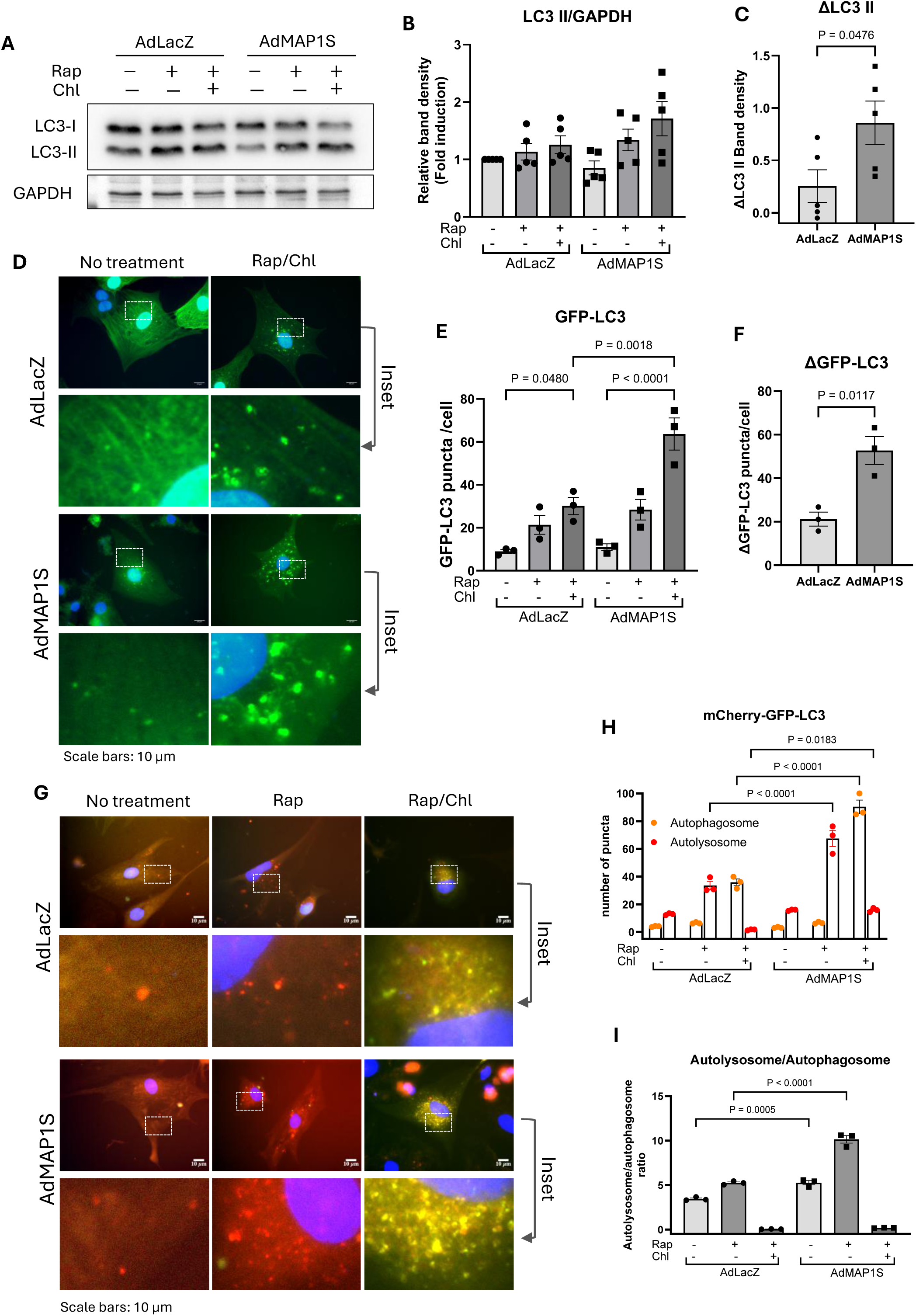
MAP1S overexpression promotes autophagic flux in cardiomyocytes. **A)** Western blots and **B)** quantification of band density of endogenous LC3-II in NRCM after rapamycin (5 μM) and chloroquine (3 μM) treatment. **C)** ΔLC3-II calculation indicated that MAP1S overexpression significantly increased the change in LC3-II levels in NRCMs following rapamycin/chloroquine treatment (n=5 independent experiments). **D) I**mages of NRCM expressing GFP-LC3 co-treated with rapamycin/chloroquine. **E)** Quantification of GFP-LC3 puncta and **F)** ΔGFP-LC3 further confirmed the increase in autophagosome accumulation after MAP1S overexpression, suggesting that MAP1S overexpression either enhanced autophagosome formation or inhibited autophagosome degradation. **G)** Images of NRCMs transduced with mCherry-GFP-LC3 and treated with rapamycin/chloroquine. **H)** Quantification of yellow and red puncta per cell. **I)** The autolysosome/autophagosome (red/yellow) ratio per cell was significantly increased in NRCM overexpressing MAP1S both basally and after rapamycin treatment, indicating increased autophagic flux. Statistical tests used: (**B, E,I**) One-way ANOVA followed by posthoc multiple comparisons. (**C,F**) Student’s t-test; (**H**) Two-way ANOVA followed by posthoc multiple comparisons. Numbers above the bars indicate *P* values. Number of replications: (**B,C**) n=5 independent experiments; (**E,F,H,I**) n=3 independent experiments.

Since higher GFP-LC3 puncta may also be interpreted as a result of autophagosome accumulation due to a reduction in autophagosome-lysosome fusion, we conducted experiments using the mCherry-GFP-LC3 construct that can differentiate the autophagosome from the autolysosome. Data presented in figures 6G-H indicated that in the presence of rapamycin, the number of autolysosomes was significantly increased in MAP1S overexpressing cells, whereas after treatment with rapamycin and chloroquine, the number of autophagosomes was elevated. Importantly, when we analyzed the autolysosome/autophagosome ratio, which is an indicator of autophagic flux, we observed a significant increase in MAP1S-expressing cells both at basal and after rapamycin treatment (Fig. 6I). Together, these data suggested that MAP1S overexpression induced autophagic flux in cardiomyocytes.

### Establishment of MAP1S overexpression using the modRNA approach

The in vitro data using the overexpression model prompted us to hypothesize that cardiac overexpression of MAP1S will be beneficial in pathological conditions such as MI. We used a modified RNA approach to express MAP1S, as it has been demonstrated as an effective way to achieve a strong transient expression of a transgene when injected intramyocardially^16^. We therefore generated modified RNA to express MAP1S and luciferase (as a control). Expressions of luciferase and MAP1S were detected in the mouse heart following direct intra-myocardial injection (Supplementary Fig.6).

### modRNA-mediated expression of MAP1S protected the heart from adverse remodeling following MI

To test the effects of MAP1S overexpression during MI, we subjected wild type C57Bl/6 mice to MI. modRNA expressing MAP1S or luciferase was injected intramyocardially straight after the coronary ligation. Cardiac phenotypes were then assessed at 1 week post-MI. There was no significant difference in heart weight/body weight ratio between the modMAP1S vs modLuc group (Fig.7A); however, analysis of cardiac function showed a significant improvement in ejection fraction and cardiac output in mice treated with modMAP1S compared to modLuc (Fig.7B-C). In addition, MAP1S overexpression reduced cardiomyocyte cell size in the MI border zone (Fig.7D-E). More importantly, apoptosis was significantly reduced in modMAP1S-treated mice (Fig.7F-G). These results were consistent with the in vitro data using the NRCM culture and the in vivo data using the MAP1S knockout model.

**Figure 7.**
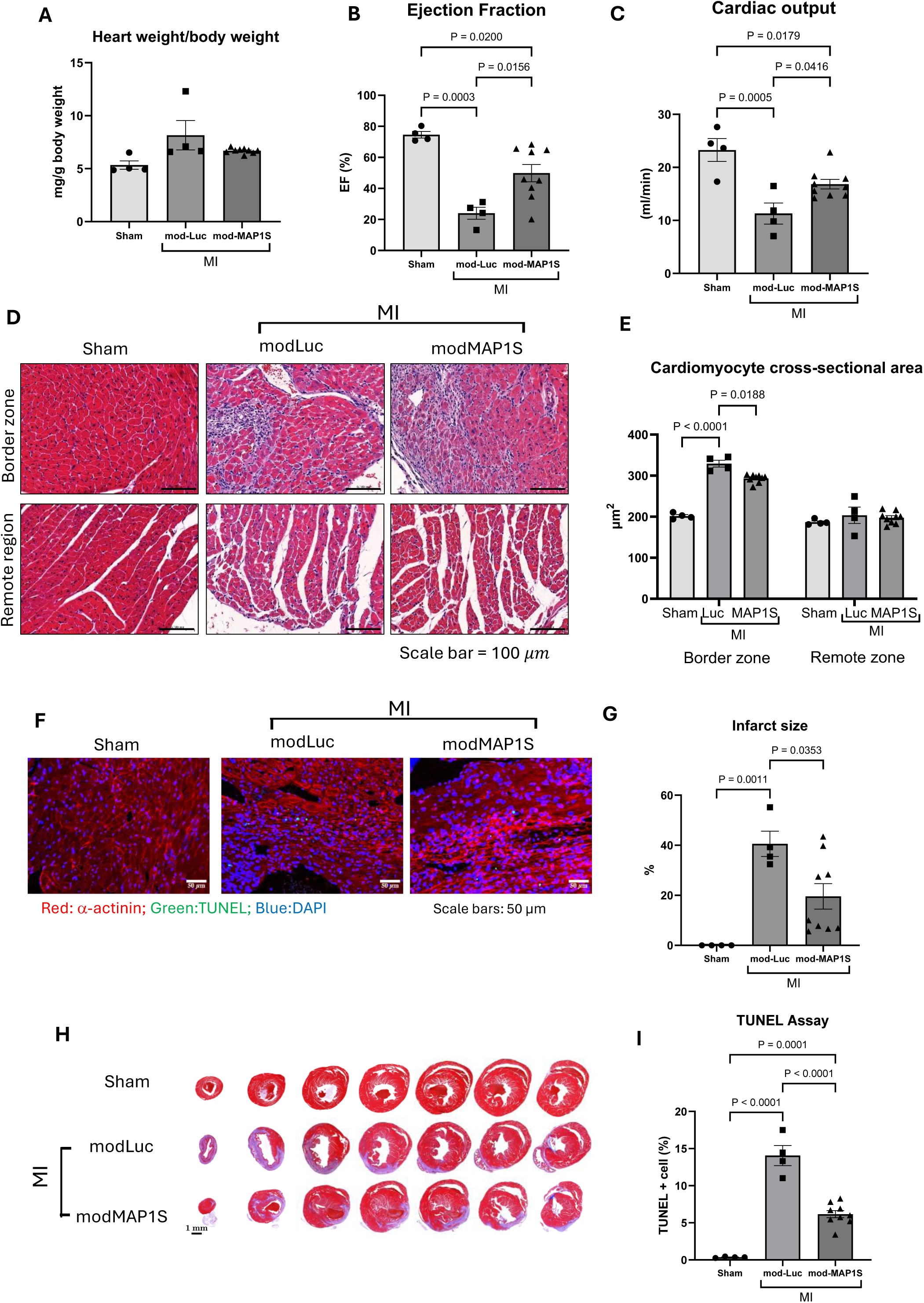
mod-RNA mediated expression of MAP1S in the heart improves cardiac function and reduces apoptosis following MI. Wild-type C57Bl/6 mice were subjected to MI and treated with intramyocardial injection of modified RNA (modRNA; 50 μg/heart) encoding either MAP1S (modMAP1S) or luciferase (modLuciferase). **A)** Analysis of heart weight/body weight ratio showed no significant difference between groups. Cardiac performance as indicated by **B)** ejection fraction and **C)** cardiac output was significantly enhanced in modMAP1S-treated mice compared to modLuciferase-treated controls 7 days after MI. **D)** H&E-stained sections and **E)** quantification of cardiomyocyte cross-sectional area showed smaller cardiomyocyte size in the infarct border zone of modMAP1S-treated mice. **F)** Representative TUNEL-stained sections followed by **G)** quantification of TUNEL-positive cells showed reduced apoptosis in modMAP1S-treated mice compared to modLuciferase-treated controls 7 days after MI. **H**) Masson’s trichrome-stained heart sections and **(I)** quantification of infarct size demonstrated a significant reduction in infarct size in modMAP1S-treated mice compared to modLuciferase-treated controls 7 days following MI. (sham, n=4; MI+modLuciferase, n=4; MI+modMAP1S, n=9). Statistical tests used: (**A,B,C,G,I)** One-way ANOVA followed by posthoc multiple comparisons; (**E**) Two-way ANOVA followed by posthoc multiple comparisons. Numbers above the bars indicate *P* values.

Infarct size was assessed using Masson-trichrome staining (Fig.7H). Analysis of the infarct area revealed that treatment with modMAP1S significantly reduced infarct size (Fig.7I). Echocardiographic analysis of cardiac morphology further strengthens our finding, as MAP1S overexpression produced a trend of better remodeling such as reduced left ventricular diameter (indicating reduced dilatation) and increased wall thickness (protection against myocardial wall thinning) compared to the control group (Supplementary figure.7A-G).

Together, our data demonstrated that transient overexpression of MAP1S was beneficial in protecting against excessive apoptosis and preserving cardiac function and structure following MI.

## DISCUSSION

The main finding of this study is that MAP1S expression exerts a protective role in cardiomyocytes and in the heart following stress. In primary cardiomyocytes, MAP1S expression increases autophagic activity and reduces apoptosis. In infarcted mouse hearts, transient overexpression of MAP1S inhibits apoptosis, reduces scar size, and improves cardiac function and morphology. Our study also provides the first evidence that MAP1S may regulate apoptosis by modulating the Hippo/YAP pathway. Therefore, MAP1S may represent a novel therapeutic target to preserve cardiomyocyte viability and mitigate the development of HF following MI.

Autophagy and apoptosis are two mechanisms heavily involved in the regulation of cardiomyocyte death and survival following MI^17^. Autophagy is a cellular process for degrading and recycling dysfunctional cytoplasmic components. It is essential for cell homeostasis and is regarded as a pro-survival response to pathological stimuli^18^. MAP1S has been demonstrated as a regulator of autophagy in non-cardiac cells, mainly through its interaction with LC3 (or Atg8), a key modulator of autophagy^11, 19^. Previous studies showed that MAP1S depletion resulted in inefficient autophagic activity in various pathological settings, including renal fibrosis^20^, liver fibrosis^19^, hepatocarcinoma^21^, clear cell renal cell carcinoma^22^, and acute promyelocytic leukaemia^23^. These studies highlight that defective autophagy due to MAP1S ablation might enhance fibrosis^19, 20^, and genome instability^21, 22^. In contrast, MAP1S overexpression resulted in autophagic induction ^21, 22, 23, 24, 25, 26^, accompanied by suppression of fibrosis^25, 26^, oxidative stress^25^, and tumorigenesis^21, 22, 25, 26, 27^. However, despite advances in the understanding of MAP1S’ functions in the setting of fibrosis and tumorigenesis, the role of MAP1S in cardiomyocytes and in the heart is less well studied.

Our findings are consistent with previous observations in non-cardiomyocytes that MAP1S functions as a positive modulator of autophagy. Ablation of MAP1S through gene silencing resulted in the accumulation of autophagosome. Autophagosome accumulation could either be due to an increase in autophagosome formation or a reduction of autophagosome degradation. Interestingly, under ischaemic condition MAP1S^-/-^ hearts displayed a reduction in lysosome/phagosome ratio, suggesting that the impairment of autophagosome degradation is more likely the cause of autophagosome accumulation in MAP1S-deficient cardiomyocytes. In addition, data from the overexpression model using the LC3-GFP-mCherry probe supports this notion.

Given that MAP1S deficiency seems to impair autophagy, we assessed if overexpression of MAP1S resulted in enhanced autophagic activity. To this end, adenovirus-mediated overexpression of MAP1S in NRCM was used as a cellular model. We used a tandem mCherry-GFP-LC3 system, which allowed us to determine the autophagic flux by simultaneous monitoring of autophagosomes and autolysosomes^28^. Results from this assay revealed that autophagy flux was increased in MAP1S overexpressing cells compared to controls. Thus, it seems that overexpression of MAP1S induces both the autophagosome formation and the autophagosome-lysosome fusion. Together, it can be postulated that MAP1S positively regulates autophagy by promoting both autophagosome biogenesis and clearance in cardiomyocytes.

Our evidence showed that deficiency of MAP1S in NRCM enhanced apoptosis in response to oxidative stress. Consistently, MAP1S gene ablation in mice led to a dramatic increase in mortality following MI with a clear sign of increased apoptosis at both the acute (3 days) and chronic (4 weeks) phases of MI. Increased apoptosis is strongly associated with cardiac rupture after MI in mice^29^. This might be the cause of the poor survival observed in MAP1S^-/-^ mice following MI. Consistently, we found that MAP1S overexpression significantly reduced cardiomyocyte apoptosis in response to oxidative stress. Expression of caspase-3, a downstream effector of both the intrinsic and extrinsic apoptosis pathways, showed a trend towards decreased activation in MAP1S-overexpressing cells.

Prior studies have suggested that MAP1S deletion caused alterations of mitochondrial morphology^11^. Indeed, mitochondria play a key role in the regulation of apoptosis by releasing pro-apoptotic factors from the mitochondrial intermembrane space^30, 31^. This process is controlled by the Bcl-2 family proteins^32^. This prompted us to speculate that MAP1S-mediated apoptosis regulation could be via its role in maintaining the mitochondria’s function and structure. However, our data indicated that in cardiomyocytes and in whole hearts, the levels of key proteins that mediate the mitochondrial apoptosis pathway (the Bcl-2 family proteins) were not altered following ablation of MAP1S. Thus, regulation of the mitochondrial pathway might not be the main mechanism responsible for the MAP1S-dependent apoptosis regulation in the heart and isolated cardiomyocytes.

We reasoned that MAP1S might regulate other apoptotic-related pathway(s) in cardiomyocytes. Interestingly, through bioinformatics analysis of MAP1S interacting partners, we found that the Hippo pathway is the most enriched pathway among MAP1S interactors. This pathway regulates numerous biological processes, including apoptosis, cell proliferation, and cell survival^33^. Notably, the serine/threonine kinases MST1 and MST2, which are the two core components of the Hippo pathway, physically interact with MAP1S in _HEK293 cells_34, 35, 36, 37, 38_._

By using isolated primary cardiomyocytes, we found that MAP1S expression reduced the phosphorylation, hence the activation of MST1/2. In contrast, MAP1S gene silencing increased phosphorylation of MST1/2. MST1/2 is known as pro-apoptotic kinases as they control apoptosis through the Hippo pathway by modulating YAP activity^39^. These kinases also regulate apoptosis via alternative pathways, such as phosphorylation of histone H2AX^40^, the FOXO3 regulation^41^ and Caspase-3-induced activation and nuclear translocation^42^. Furthermore, our data suggests that the activity of YAP, which is the main downstream Hippo pathway effector, is modulated by MAP1S expression. YAP nuclear translocation and co-transcriptional activity were enhanced following MAP1S overexpression and reduced after gene silencing. It is known that YAP activation leads to the expression of anti-apoptotic factors and pro-survival proteins, such as Bcl2, Bcl2l2, and survivin^43, 44^. Therefore, our findings indicate that MAP1S possibly regulates cardiomyocyte apoptosis in cardiomyocytes by modulation of the Hippo/YAP pathway.

Perhaps the most interesting finding from the translational point of view is the result that modRNA-mediated expression of MAP1S protects the heart from functional and structural damages following MI. MAP1S overexpression resulted in attenuated myocardial apoptosis, reduction of infarct size, and LV wall thinning. It also improves contractile functions, such as ejection fraction and cardiac output. modRNA-mediated gene expression has recently emerged as a potential approach to induce expression of a therapeutic gene as an alternative to a viral-based system. This technique allows rapid and transient gene expression, making it an ideal tool to introduce protective genes soon after MI to prevent cardiomyocyte loss. It represents a safer option compared to other gene delivery systems because it lacks immunogenicity and does not integrate into the host genome, which prevents unwanted long-term effects^45^.

Despite encouraging results from overexpressing MAP1S in primary cardiomyocytes and in the mouse heart, there are still several questions that need addressing. For example, the possible connection between the regulation of autophagy and apoptosis by MAP1S remains unclear. It would be interesting to understand whether MAP1S provides autophagic protection against apoptosis induction or whether MAP1S regulates autophagy and apoptosis in an independent manner. Future studies should also explore the effects of MAP1S expression in other pathological conditions, such as pressure overload and ischaemia-reperfusion injury. It is also important to understand the role of MAP1S in other cell types, such as cardiac fibroblasts and residential macrophages, which play essential roles in cardiac remodeling. This is important because in the modRNA system we did not use a cardiomyocyte-specific targeting approach, and so the effects on fibroblasts and macrophages may contribute to the phenotype.

We acknowledge that this study has several limitations. First, we performed permanent coronary ligation in mice, which models MI without reperfusion relevant to ∼25-30% of clinical MI^46^. However, since 70–75% of clinical MI now involve reperfusion therapy^46^, it is important to investigate the role of MAP1S in ischaemia/reperfusion models in future studies. Second, the use of global MAP1S knockout prevents us from determining if MI-induced mortality stems solely from cardiac pathologies or involves other organs. Also, this model did not provide the cell-specific role of MAP1S. Future studies using tissue- or cell-specific knockout models will be essential to clarify these systemic effects and identify cell-specific signalling pathways.

In conclusion, our work demonstrates that MAP1S exerts a protective effect in cardiomyocytes and in the heart by inhibiting apoptosis following pathological stimulation. This phenotype might be associated with its interaction and regulation of the Hippo/YAP pathway. In addition, positive modulation of autophagy by MAP1S may also contribute to this protective function in the heart and cardiomyocytes. We also demonstrate the beneficial effects of modRNA-mediated MAP1S expression in a therapeutic model of MI in mice, suggesting that MAP1S may represent a novel therapeutic target to preserve cardiomyocyte viability and mitigate HF.

## METHODS

### Analysis of MAP1S expression

Heart tissue extracts were collected from C57Bl/6 mice (8 weeks old) following permanent coronary artery ligation (myocardial infarction/MI) for 4 weeks. Details of MI protocols are in Supplementary methods. MAP1S expression was detected using Western blot as detailed in Supplementary methods. We analysed datasets available in the Gene Expression Omnibus (GEO) database: GSE163956^47^ and GSE116250^48^ for studying MAP1S cell specific expression during MI (GSE163956) and MAP1S expression in human heart tissues (GSE116250). GSE116250 dataset includes RNA sequencing data of heart tissues from patients with heart failure with ischaemic cardiomyopathy (HF-ICM), heart failure with dilated cardiomyopathy (HF-DCM) and non-failing hearts. Detailed patients’ information are available in previous publications^48^. Data were analysed using GEO2R analysis tool to determine differentially expressed genes. Please see Supplementary methods for analysis of MAP1S cell specific expression using GSE163956 dataset.

### Isolation and culture of Neonatal rat cardiomyocytes and cardiac fibroblasts

Neonatal rat cardiomyocytes (NRCMs) and cardiac fibroblasts (NRCF) were isolated from 2-3-day-old Sprague-Dawley rat neonates’ hearts using enzymatic dissociation as previously described^49^. Please see Supplemental Materials for detailed protocols on cell isolation and culture, MAP1S gene silencing and overexpression in cellular model, and analysis of cellular phenotypes including apoptosis, autophagic flux, mitochondrial function and Hippo pathway/YAP activity assays.

### Animal studies

Mice were housed in the University of Manchester BSF Facility in a standard laboratory animal housing condition. Mice were maintained on a 12-hour light/dark cycle in a controlled temperature of 19-22°C and humidity of 40-65%. Mice were fed with standard chow diet (BK001, Special Diet Services, UK) and water ad libitum. Mice with systemic deletion of *Map1s* gene (MAP1S^-/-^ mice) were used in this study. The generation of these knockout mice was described elsewhere^11^. These mice were maintained in C57Bl/6 background. Genotyping was performed using standard PCR procedure using primers as described in Supplementary Table 1. Wild-type C57BL/6J mice were also used in this study. Twelve-week-old C57BL/6J mice were purchased from Envigo UK and were allowed to acclimate for a week before in vivo experiments were performed.

Detailed methods on MI model, phenotypic characterization following MI and modRNA-mediated cardiac overexpression are available in Supplemental Materials.

### Ethical declaration

All animal experiments were carried out in accordance with the United Kingdom Animals (Scientific Procedures) Act 1986 under Home Office Project Licence PP5982529. All procedures were reviewed and approved by the University of Manchester Animal Welfare and Ethical Review Board. All animal experiments were performed and is reported in compliance with the ARRIVE (Animal Research: Reporting of In Vivo Experiments) guidelines.

### Statistical analysis

Statistical analysis was conducted using one- or two-way ANOVA followed by Tukey’s post hoc tests where appropriate, and comparisons between two groups were performed using Student’s t-test. Survival analysis was carried out using the Log-rank (Mantel-Cox) test. All data are presented as mean ± SEM. We used GraphPad Prism software ver.10.2.2 (GraphPad Software, LLC) for statistical analysis and data presentation. A p-values < 0.05 were considered statistically significant.

## DATA AVAILABILITY

All data generated in this study are presented in Results section and in Supplemental Materials. Data are available from the corresponding author upon reasonable request.

## ACKNOWLEDGEMENTS

We thank the University of Manchester Biological Services Facility and the Bioimaging Facilities for technical support. The authors thank the staff in the Electron Microscopy Core Facility in the Faculty of Biology, Medicine and Health, University of Manchester for their assistance, and the Wellcome Trust for equipment grant support to the EM Core Facility

## AUTHOR CONTRIBUTIONS

P.M. planned, designed and performed in vivo and in vitro experiments, analysed data, conducted bioinformatics studies and wrote manuscript. Y.S.K. performed studies on MAP1S^-/-^ mice. S.P. and M.Z. performed in vivo surgical experiments. A.B.N. and E.T. helped in conducting in vivo and in vitro experiments. D.P. conducted electron microscopy analysis. G.G. supervised and helped mitochondria function analysis. E.J.C. supervised and helped designing in vivo experiments. L.L. provided reagents and helped with data interpretation. D.O. conceived the scientific ideas, oversaw the project, designed in vivo and in vitro experiments, analysed data and wrote the manuscript.

