## Supplemental Materials for "Microtubule-associated protein 1S (MAP1S): a cardioprotective factor against post-myocardial infarction remodeling via apoptosis inhibition"

**Short title:** Role of MAP1S in regulating heart remodeling

5.002, AV Hill Building, The University of Manchester
Oxford Road, Manchester M13 9PT, United Kingdom

<sup>†</sup>Deceased, 11 February 2024

**Supplementary Figure 1 - 7**

**Supplementary Table 1 – 4**

**Supplementary Methods**

**References**

**Supplementary Figure 1**

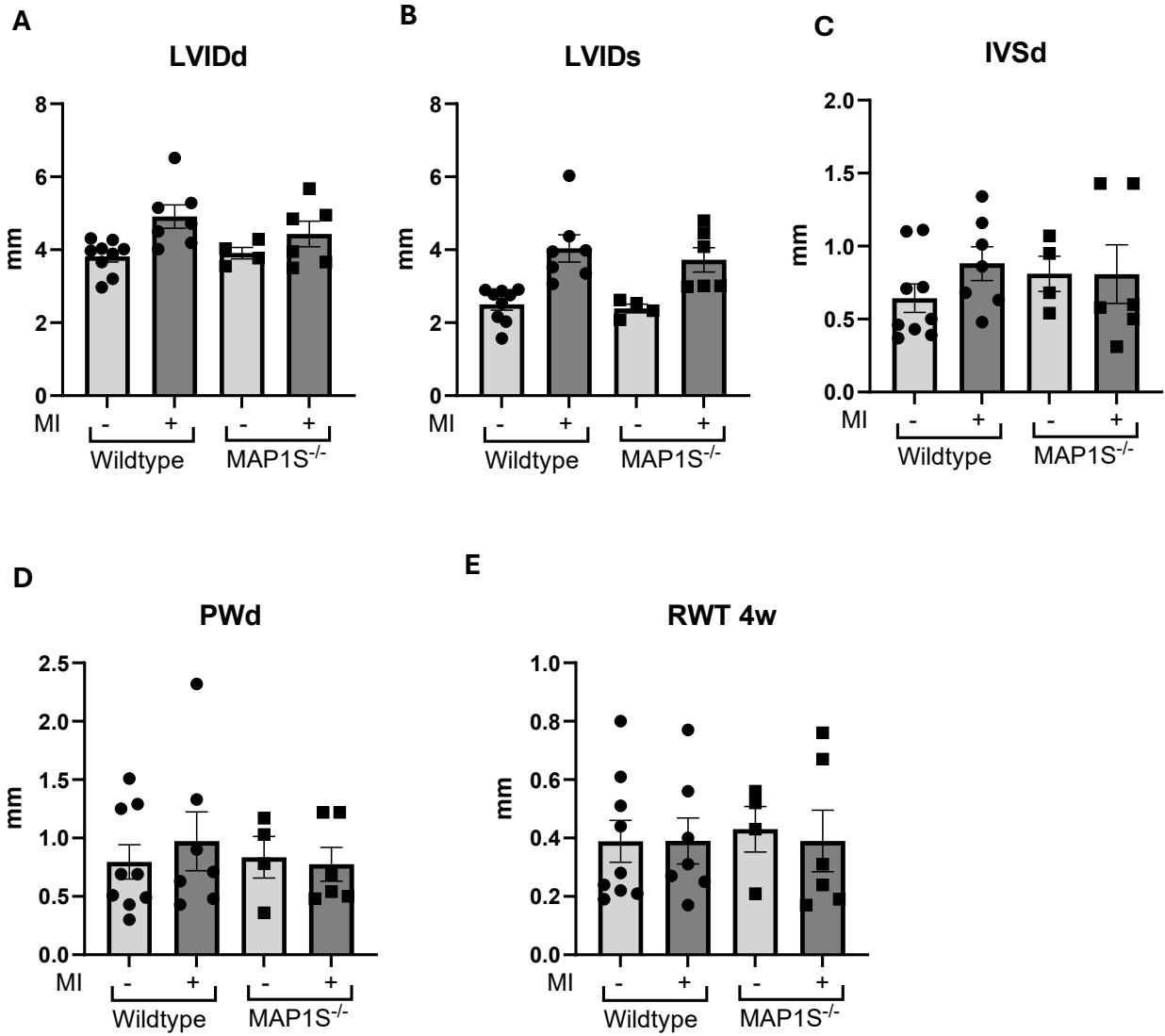

**Supplementary figure 1 Echocardiography analysis of MAP1S<sup>-/-</sup> and WT littermates at 4 weeks after MI.**

Echocardiography analysis showed that **A)** left ventricular internal diameter at diastole (LVID<sub>d</sub>), **B)** left ventricular internal diameter at systole (LVID<sub>s</sub>), **C)** interventricular septal wall thickness at diastole (IVS<sub>d</sub>), **D)** posterior wall at diastole (PW<sub>d</sub>), and **E)** relative wall thickness were not significantly different between MAP1S<sup>-/-</sup> and WT littermates. (WT sham, n=9; WT MI, n=7; MAP1S<sup>-/-</sup> sham, n=4; MAP1S<sup>-/-</sup> MI, n=6). Data are presented as mean ± SEM.

### Supplementary Figure 2

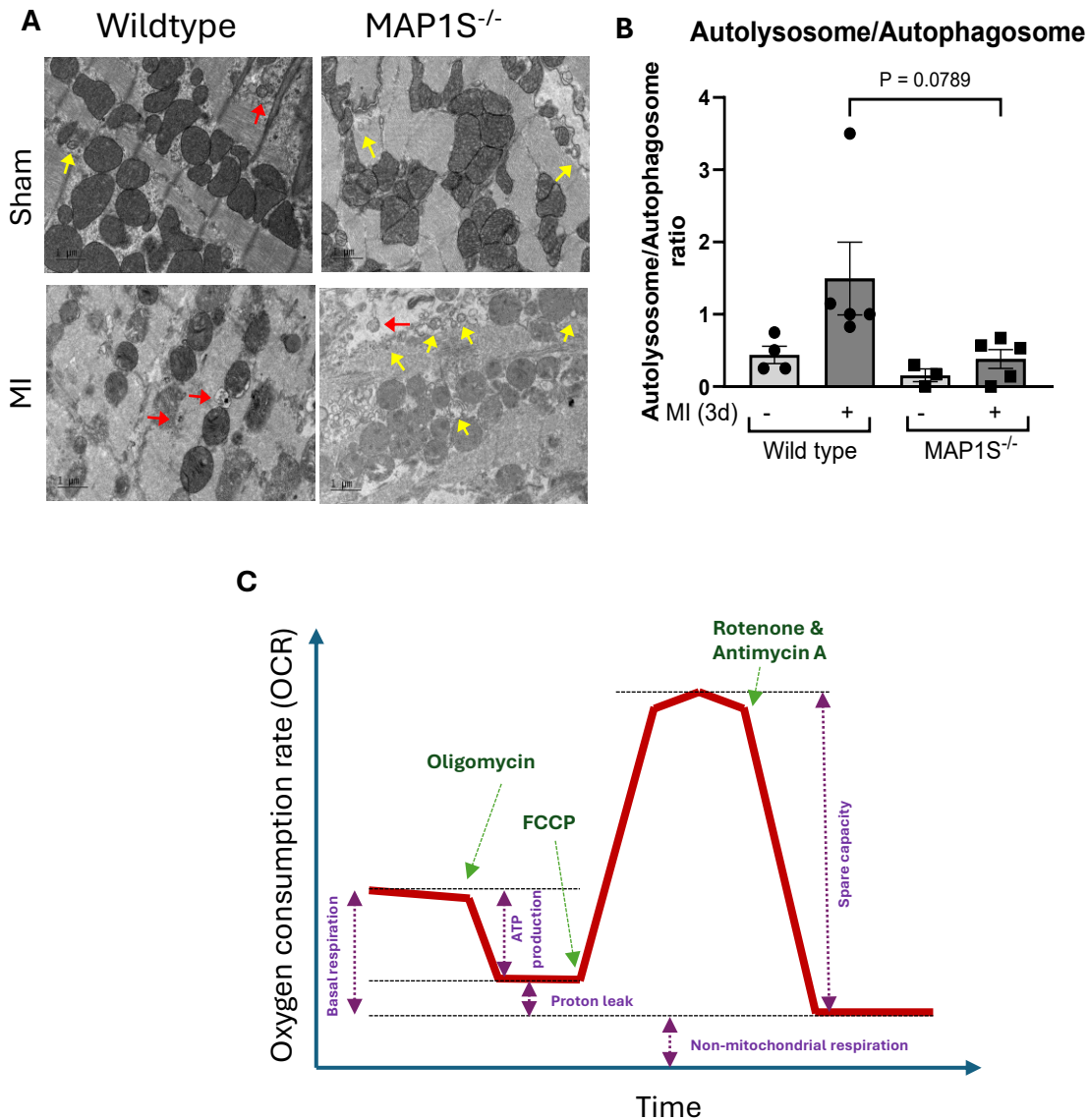

**Supplementary Figure 2** MAP1S<sup>-/-</sup> mice and wild-type (WT) littermates were subjected to MI. Hearts were harvested 3 days after MI for transmission electron microscopy (TEM) analysis. **A)** Representative TEM images showing autophagosomes (yellow arrow) and autolysosomes (red arrow) (scale bars = 1 μm). **B)** The autolysosome/autophagosome ratio, an index of autophagic flux, was calculated based on the number of autophagic vesicles in MAP1S<sup>-/-</sup> and WT hearts. (WT sham, n=4; WT MI, n=5; MAP1S<sup>-/-</sup> sham, n=3; MAP1S<sup>-/-</sup> MI, n=5). **C)** Schematic diagram depicting the Seahorse Mito Stress Test to measure oxygen consumption in cultured cells. The oxygen consumption rate (OCR) is calculated in the presence of mitochondria respiration inhibitors to assess specific aspects of mitochondria function, including basal respiration, ATP-linked respiration, maximal respiration and reserve respiratory capacity (see Fig.4).

#### Supplementary Figure 3

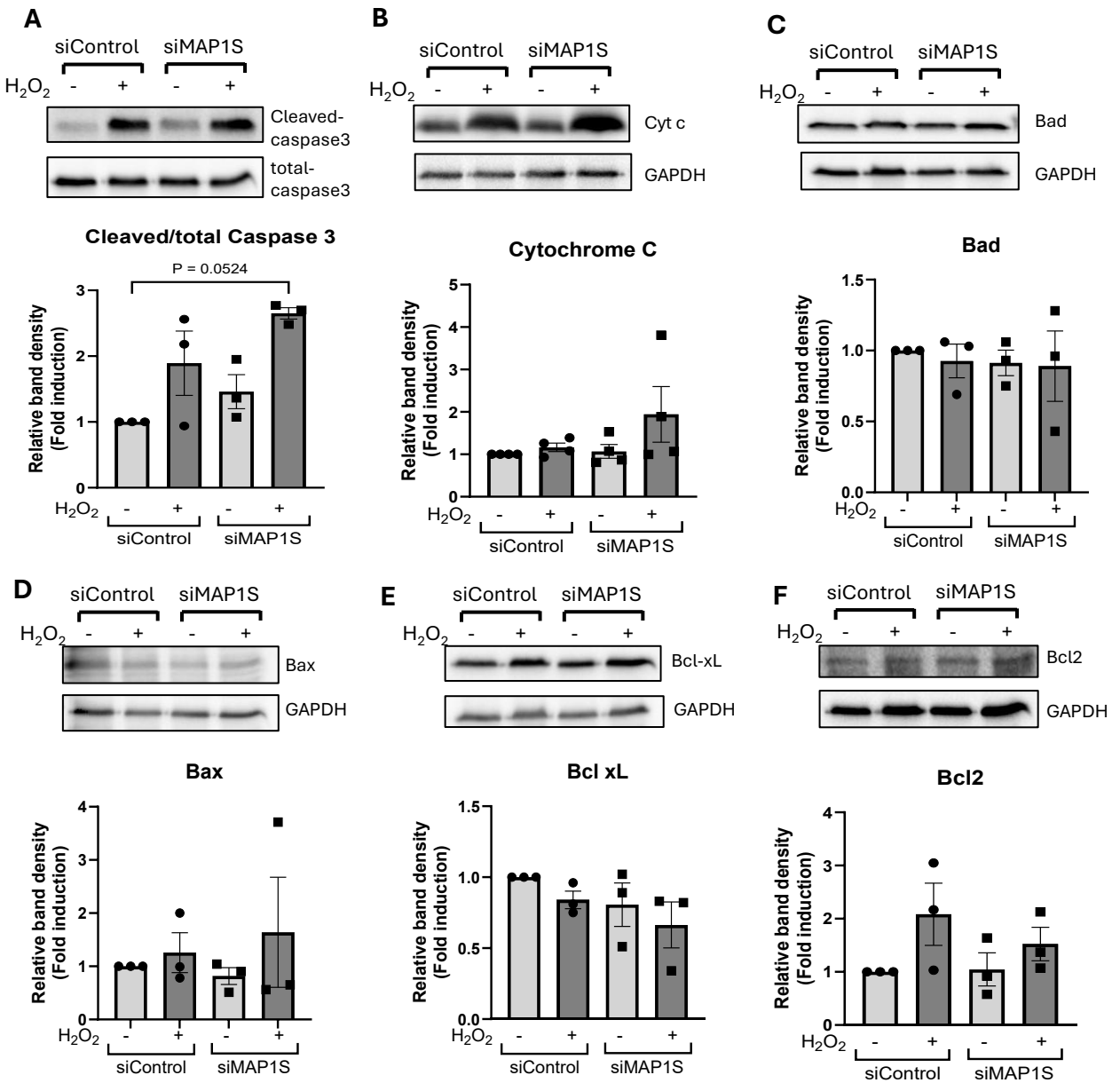

#### Supplementary figure 3 Expressions of mitochondria-related apoptosis regulators in neonatal rat cardiomyocytes (NRCM) in response to MAP1S gene silencing.

Western blot analysis was performed to assess the expression of apoptosis regulators in MAP1S-depleted NRCMs following oxidative stress. **A)** Western blots showing expressions of cleaved and total caspase 3 in MAP1S-depleted NRCMs treated with H<sub>2</sub>O<sub>2</sub> (100 μM for 2 hours). The cleaved/total caspase-3 ratio showed a trend toward increased caspase-3 activation (n = 3 independent experiments). However, immunoblot analysis of mitochondria-related apoptosis regulators: **B)** cytochrome c, **C)** Bad, **D)** Bax, **E)** Bcl-xL, and **F)** Bcl-2 showed no significant differences between MAP1S-depleted NRCMs and control cells following H<sub>2</sub>O<sub>2</sub>-induced oxidative stress (n = 3-4 independent experiments).

### Supplementary Figure 4

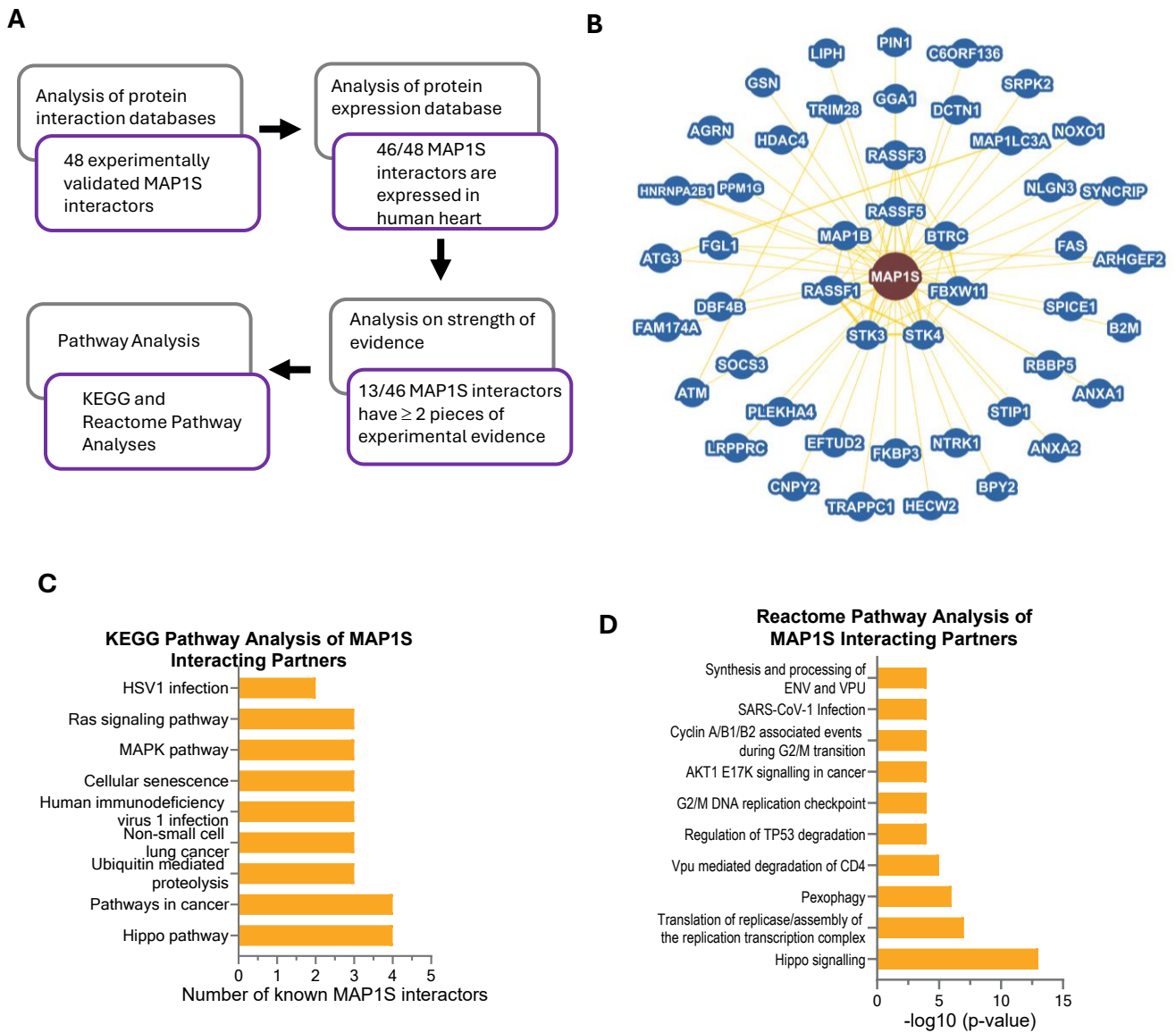

**Supplementary Figure 4 Analysis of MAP1S interacting proteins.** A) Scheme describing the process of the analysis of MAP1S interacting proteins. List of interacting partners are shown in supplementary table 1. B) MAP1S protein interaction network visualised by BioGRID. C) KEGG pathway analysis and D) Reactome overrepresentation analysis both revealed that the Hippo pathway is the most enriched pathway among MAP1S-interacting partners.

**Supplementary Figure 5**

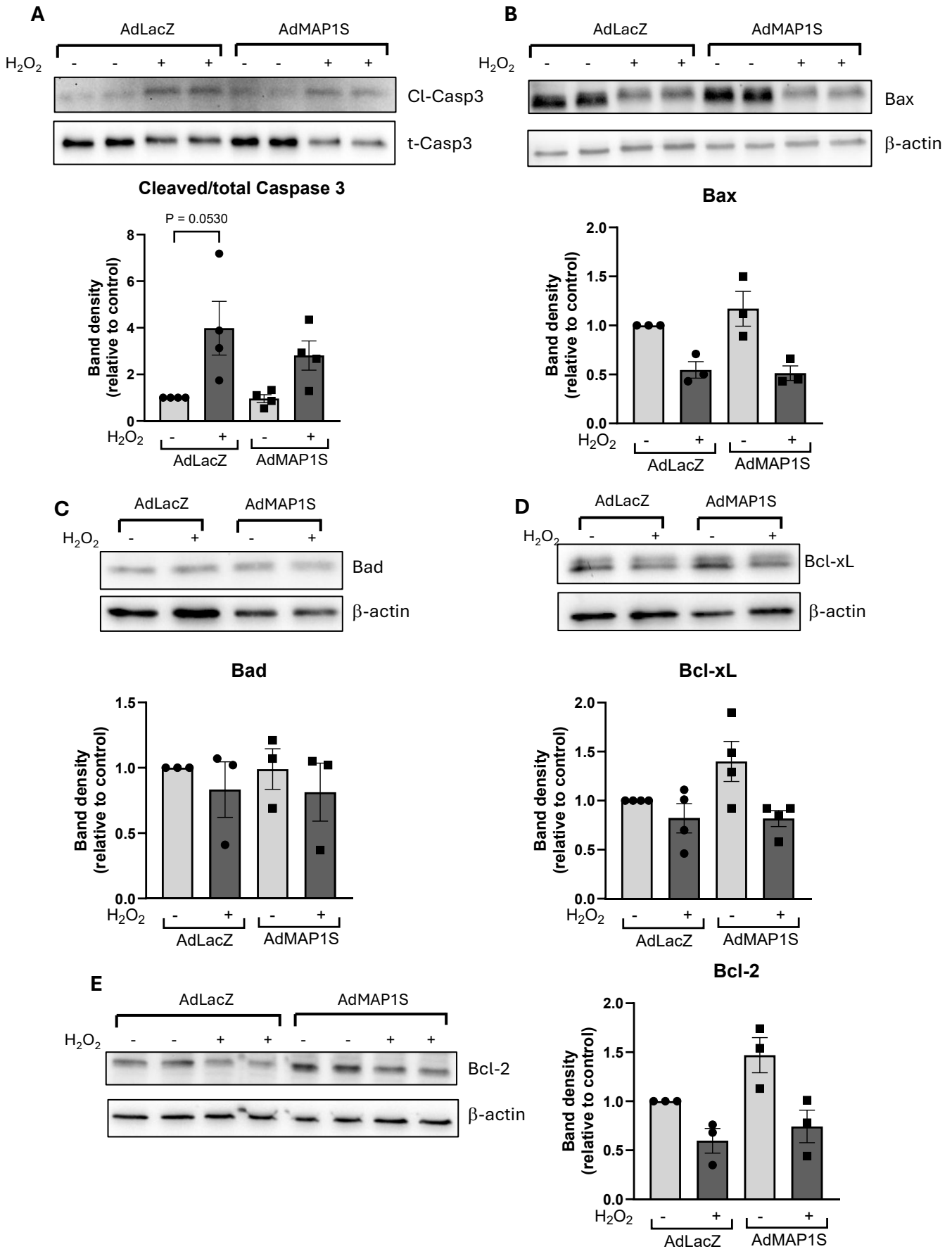

**Supplementary figure 5 MAP1S overexpression does not affect expressions of mitochondria-related apoptosis mediators in cardiomyocytes.**

**A)** Examples of Western blot images showing expression of cleaved and total caspase 3 in MAP1S-overexpressing cells treated with H<sub>2</sub>O<sub>2</sub> (100 μM for 2 hours). The cleaved/total caspase-3 ratio showed a trend toward reduced caspase-3 activation in MAP1S-overexpressing NRCMs compared to control cells (n = 3 independent experiments). However, immunoblot analysis of mitochondrial-related apoptosis markers **B)** Bax, **C)** Bad, **D)** Bcl-xL, and **E)** Bcl-2 showed no significant differences between MAP1S-overexpressing and control NRCMs following H<sub>2</sub>O<sub>2</sub>-induced oxidative stress (n = 3-4 independent experiments using different batches of NRCMs).

### Supplementary Figure 6

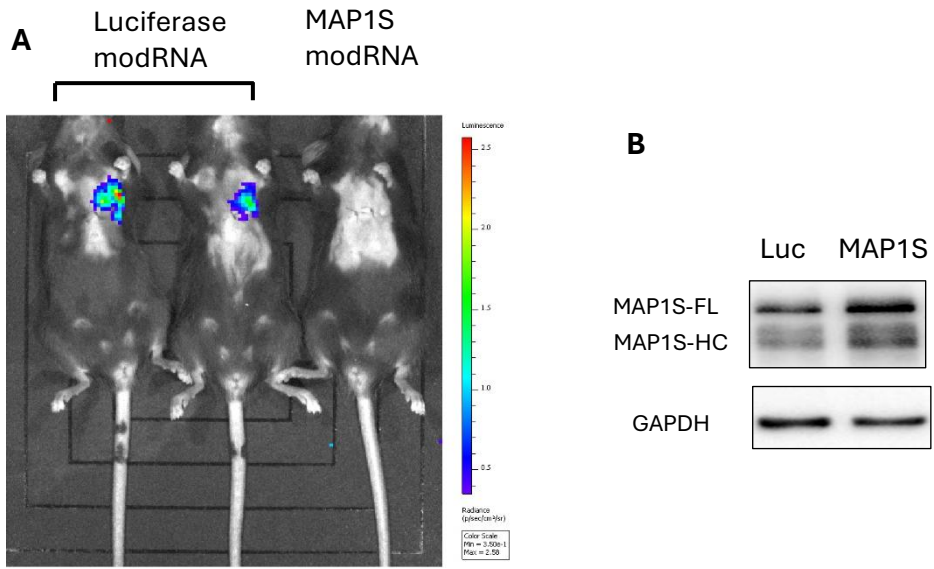

#### Supplementary figure 6 Validation of mod-RNA mediated expression of luciferase and MAP1S in mouse hearts.

Luciferase or MAP1S modRNA were injected intramyocardially in WT C57Bl/6 mice (50 μg/heart). **A)** Luciferase modRNA yielded a robust bioluminescent signal 24 hours after injection, indicating localised luciferase protein expression in the heart. **B)** Western blot showing that modRNA-mediated delivery of the MAP1S gene efficiently increased MAP1S protein expression 48 hours after intramyocardial injection.

### Supplementary Figure 7

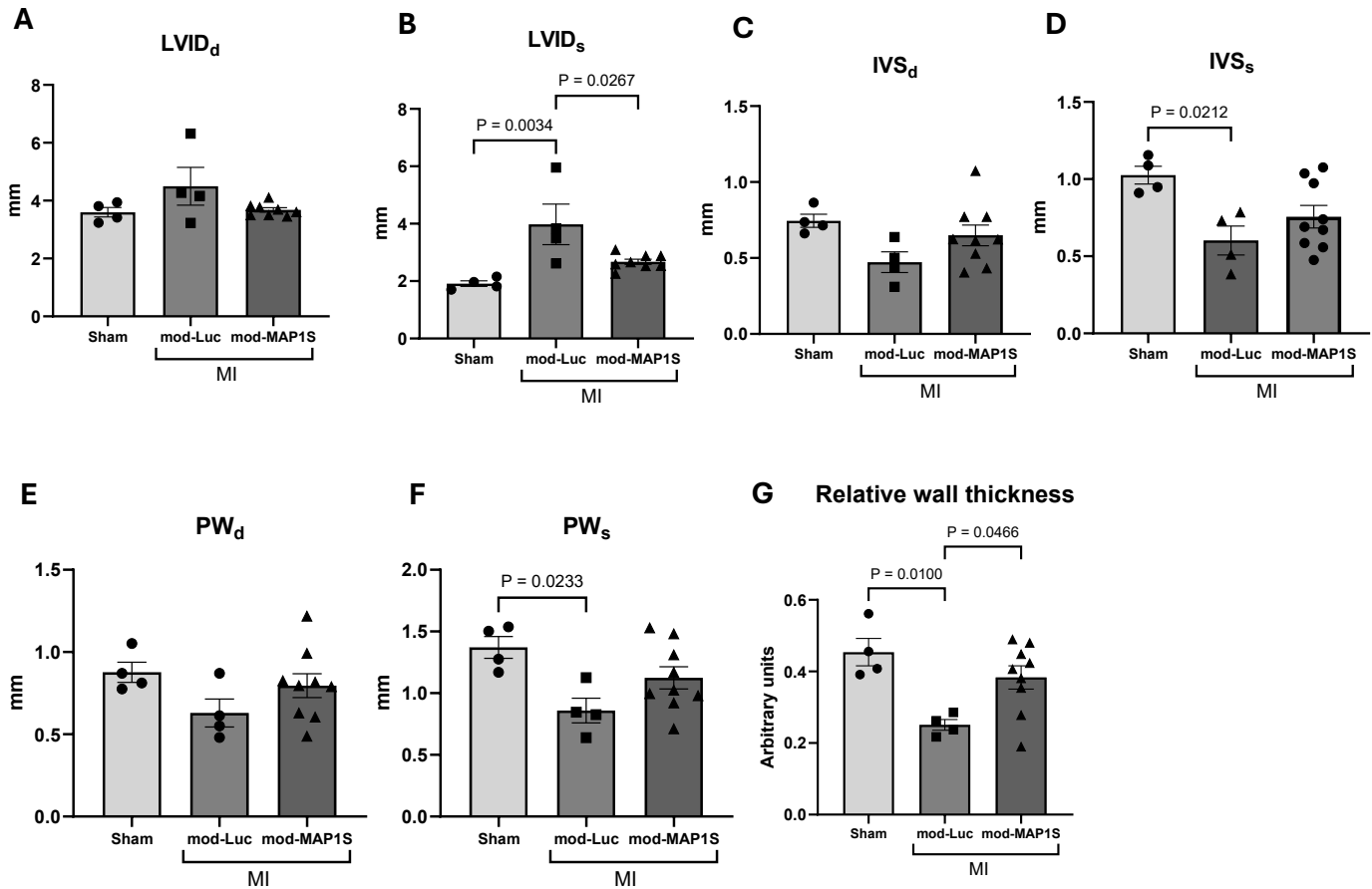

#### Supplementary Figure 7 Echocardiography analysis indicates that mod-RNA mediated expression of MAP1S improves cardiac remodelling after MI.

**A)** Left ventricular internal diameter at diastole (LVID<sub>d</sub>) did not differ between groups. However, **B)** LVID at systole (LVID<sub>s</sub>) was significantly reduced in modMAP1S-treated mice, indicating attenuated ventricular dilatation. **C)** Septal wall thickness at diastole (IVS<sub>d</sub>) was comparable between groups. **D)** Septal wall thickness at systole (IVS<sub>s</sub>) was significantly reduced in modLuciferase-treated mice, but remained preserved in modMAP1S-treated mice. **E)** Posterior wall thickness at diastole (PW<sub>d</sub>) did not differ between groups. **F)** Posterior wall thickness at systole (PW<sub>s</sub>) was significantly reduced in modLuciferase-treated mice, but remained preserved in modMAP1S-treated mice. **G)** Consistently, the relative wall thickness was significantly greater in modMAP1S-treated hearts compared to modLuciferase-treated controls, indicating protection against left ventricular wall thinning. (sham, n=4; MI+modLuciferase, n=4; MI+modMAP1S, n=9).

**Supplementary Table 1. Primers sequences and size of PCR products for the genotyping of MAP1S<sup>-/-</sup> mice.**

| Primer | Sequence (5' -3') | Amplicon size (bp) |  |  |
| --- | --- | --- | --- | --- |
|  |  | WT allele | KO allele | HT alleles |
| P-31 forward | CACCTGCCTAAGCCATCTGTGTC | 200 | — | 200 |
| P-32 reverse | CTCAGTCTGTCTGAGACAAGGTC |  |  |  |
| PNeo forward | GGTAGAATTGGTCGAGGTCGAC | — | 400 | 400 |
| P-32 reverse | CTCAGTCTGTCTGAGACAAGGTC |  |  |  |

WT, wild type; KO, knockout; HT, heterozygous.

**Supplementary Table 2 List of MAP1S interacting partners**

| Gene Symbol | Interaction detection method | Source database | Number of experimental evidence | Is it expressed in the human heart? |
| --- | --- | --- | --- | --- |
| RASSF1 | Affinity capture-MS/<br>affinity capture-Western/<br>two-hybrid | BioGRID/ HitPredict/<br>STRING/ IntAct/<br>Mentha/ WiKi-Pi/<br>NCBI | 6 | Yes |
| LRPPRC | Affinity capture-Western/ reconstituted<br>complex/ two-hybrid | BioGRID/ HitPredict/<br>Mentha/ WiKi-Pi/<br>NCBI | 3 | Yes |
| STK3 | Affinity capture-MS/<br>proximity label-MS | BioGRID/ HitPredict/<br>STRING/ IntAct/<br>Mentha/ NCBI | 3 | Yes |
| STK4 | Affinity capture-MS/<br>proximity label-MS | BioGRID/ HitPredict/<br>STRING/ IntAct/<br>Mentha/ NCBI | 3 | Yes |
| RASSF5 | Affinity capture-MS/<br>two-hybrid | BioGRID/ HitPredict/<br>IntAct/ Mentha/ WiKi-Pi/ NCBI | 3 | Yes |
| FBXW11 | Affinity capture-MS | BioGRID/ HitPredict/<br>Mentha/ NCBI | 3 | Yes |
| BTRC | Affinity capture-MS/<br>proximity label-MS | BioGRID/ HitPredict/<br>IntAct/ Mentha/ NCBI | 3 | Yes |
| LIPH | Affinity capture-MS | BioGRID/ HitPredict/<br>IntAct/ Mentha/ NCBI | 3 | Yes |
| MAP1LC3A | Affinity capture-Western/ reconstituted<br>complex | BioGRID/ HitPredict/<br>STRING/ IntAct/<br>Mentha/ WiKi-Pi/<br>NCBI | 2 | Yes |
| DBF4B | Affinity capture-MS | BioGRID/ HitPredict/<br>IntAct/ Mentha/ NCBI | 2 | Yes |
| FAS | Affinity capture-MS | BioGRID/ HitPredict/<br>IntAct/ Mentha/ NCBI | 2 | Yes |
| SOCS3 | Affinity capture-Western/ two-hybrid | BioGRID/ HitPredict/<br>STRING/ IntAct/<br>Mentha/ WiKi-Pi/<br>NCBI | 2 | Yes |
| BPY2 | Reconstituted complex/<br>two-hybrid | BioGRID/ HitPredict/<br>IntAct/ Mentha/ WiKi-Pi/ NCBI | 2 | No |
| FGL1 | Affinity capture-MS | BioGRID/ HitPredict/<br>IntAct/ Mentha/ NCBI | 2 | Yes |

| Gene Symbol | Interaction detection method | Source database | Number of experimental evidence | Expression in human heart |
| --- | --- | --- | --- | --- |
| ATM | Two-hybrid | BioGRID/ HitPredict/ IntAct/ Mentha/ NCBI | 1 | Yes |
| RASSF3 | Affinity chromatography technology/ anti-tag coimmunoprecipitation | BioGRID/ HitPredict/ STRING/ IntAct/ Mentha/ NCBI | 1 | Yes |
| DCTN1 | Proximity label-MS | BioGRID/ HitPredict/ IntAct/ Mentha/ NCBI | 1 | Yes |
| MAP1B | Affinity capture-MS | BioGRID/ HitPredict/ IntAct/ Mentha/ NCBI | 1 | Yes |
| STIP1 | Affinity capture-MS | BioGRID/ HitPredict/ IntAct/ Mentha/ NCBI | 1 | Yes |
| ANXA2 | Affinity capture-MS | BioGRID/ HitPredict/ IntAct/ Mentha/ NCBI | 1 | Yes |
| ANXA1 | Affinity capture-MS | BioGRID/ HitPredict/ IntAct/ NCBI | 1 | Yes |
| PPM1G | Affinity capture-MS | BioGRID/ HitPredict/ IntAct/ Mentha/ NCBI | 1 | Yes |
| SPICE | Proximity label-MS | BioGRID/ HitPredict/ IntAct/ Mentha/ NCBI | 1 | Yes |
| B2M | Affinity capture-MS | BioGRID/ HitPredict/ IntAct/ Mentha/ NCBI | 1 | Yes |
| GSN | Affinity capture-MS | BioGRID/ HitPredict/ IntAct/ Mentha/ NCBI | 1 | Yes |
| NLGN3 | Two-hybrid | BioGRID/ HitPredict/ IntAct/ Mentha/ NCBI | 1 | Yes |
| PIN1 | Reconstituted complex | BioGRID/ HitPredict/ Mentha/ WiKi-Pi/ NCBI | 1 | Yes |
| SRPK2 | Biochemical activity | BioGRID/ HitPredict/ Mentha/ NCBI | 1 | Yes |
| HDAC4 | Two-hybrid | BioGRID/ HitPredict/ Mentha/ WiKi-Pi/ NCBI | 1 | Yes |
| GGA1 | Affinity capture-MS | BioGRID/ HitPredict/ IntAct/ Mentha/ NCBI | 1 | Yes |
| ATG3 | Affinity capture-MS | BioGRID/ HitPredict/ IntAct/ Mentha/ NCBI | 1 | Yes |
| ARHGEF2 | Proximity label-MS | BioGRID/ HitPredict/ NCBI | 1 | Yes |
| EFTUD2 | Affinity capture-MS | BioGRID/ HitPredict/ NCBI | 1 | Yes |
| NTRK1 | Affinity capture-MS | BioGRID/ HitPredict/ Mentha/ NCBI | 1 | Yes |
| TRAPPC1 | Affinity capture-MS | BioGRID/ HitPredict/ IntAct/ Mentha/ NCBI | 1 | Yes |
| HECW2 | Affinity capture-MS | BioGRID/ HitPredict/ Mentha/ NCBI | 1 | Yes |
| MAP1LC3B | Anti tag coimmunoprecipitation | HitPredict/ IntAct/ Mentha | 1 | Yes |
| FAM174A | Affinity capture-MS | BioGRID/ HitPredict/ Mentha/ NCBI | 1 | Yes |
| PLEKHA4 | Affinity capture-MS | BioGRID/ NCBI | 1 | Yes |
| ATAT1 | Affinity capture-MS | BioGRID | 1 | Yes |
| NOXO1 | Affinity capture-MS | BioGRID | 1 | No |
| AGRN | Affinity capture-MS | BioGRID/ NCBI | 1 | Yes |
| TRIM28 | Affinity capture-MS | BioGRID/ IntAct/ NCBI | 1 | Yes |
| HNRNPA2B1 | Co-fractionation | BioGRID/ Mentha/ NCBI | 1 | Yes |
| FKBP3 | Co-fractionation | BioGRID/ Mentha/ NCBI | 1 | Yes |
| SYNCRIP | Co-fractionation | BioGRID/ Mentha/ NCBI | 1 | Yes |
| CNPY2 | Co-fractionation | BioGRID/ STRING/ Mentha/ NCBI | 1 | Yes |
| RBBP5 | Affinity capture-MS | BioGRID/ NCBI | 1 | Yes |

**Supplementary Table 3. List of antibodies used in this study.**

| **Antibodies** | **Species** | **Source** | **Dilution** |
| --- | --- | --- | --- |
| **Western blot** |  |  |  |
| MAP1S | Mouse monoclonal | Precision Antibody (#AG10006) | 1:1000 |
| LC3B | Rabbit polyclonal | Novus Biologicals (#NB100-2220) | 1:1000 |
| Cleaved caspase 3 (Asp175) | Rabbit polyclonal | Cell Signaling Technology (#9661) | 1:500 |
| Caspase 3 | Rabbit monoclonal | Cell Signaling Technology (#9662) | 1:1000 |
| Bax | Mouse monoclonal | Santa Cruz Biotechnology (#sc-7480) | 1:1000 |
| Bad | Rabbit polyclonal | Cell Signaling Technology (#9292) | 1:1000 |
| Bcl-xL (54H6) | Rabbit monoclonal | Cell Signaling Technology (#2764) | 1:1000 |
| Bcl-2 (D17C4) | Rabbit monoclonal | Cell Signaling Technology (#3498) | 1:1000 |
| phospho MST1 (Thr183)/MST2 (Thr180) (E7U1D) | Rabbit polyclonal | Cell Signaling Technology (#49332) | 1:500 |
| MST1 | Rabbit monoclonal | Cell Signaling Technology (#3682) | 1:1000 |
| Anti-active YAP1 (EPR19812) | Rabbit monoclonal | Abcam (#ab2052270) | 1:1000 |
| YAP | Mouse monoclonal | Santa Cruz Biotechnology (#sc-101199) | 1:500 |
| GAPDH (HRP-linked) | Rabbit | Cell Signaling Technology (#3683) | 1:5000 |
| β-actin (HRP-linked) | Rabbit | Cell Signaling Technology (#5125) | 1:5000 |
| Anti-rabbit IgG (HRP-linked) | Rabbit | Cell Signaling Technology (#7074) | 1:5000 |
| Anti-mouse IgG (HRP-linked) | Mouse | Cell Signaling Technology (#7076) | 1:5000 |
| **Immunofluorescence** |  |  |  |
| Sarcomeric α-actinin | Mouse monoclonal | Sigma-Aldrich (#A7811) | 1:200 |
| Alexa fluor 647 goat anti-mouse IgG F (ab')<sub>2</sub> fragment specific | Mouse polyclonal | Jackson Immuno Research (#115-605-072) | 1:200 |

**Supplementary Table 4. Composition of the in vitro transcription reaction for luciferase and MAP1S modRNA.**

|  | **Source** | **Concentration** | **Volume (µl)** |
| --- | --- | --- | --- |
| ARCA | Stratech | 10 mM | 3.33 |
| GTP | Thermo Fisher Scientific | 2.7 mM | 0.72 |
| ATP | Thermo Fisher Scientific | 8.1 mM | 2.16 |
| CTP | Thermo Fisher Scientific | 8.1 mM | 2.16 |
| N<sup>1</sup>-Methylpseudo-UTP | Jena Bioscience | 2.7 mM | 0.72 |
| Nuclease-free water | Thermo Fisher Scientific |  | 10.91 |
| 10X reaction buffer | Thermo Fisher Scientific |  | 2.00 |
| Enzyme mix | Thermo Fisher Scientific |  | 2.00 |
| Linearised plasmid |  | 50 ng/ µl | 1.00 |
|  |  | Total volume | 20.00 |

ARCA, anti-reverse cap analogue.

### **SUPPLEMENTARY METHODS**

#### **Analysis of MAP1S cell specific expression during myocardial infarction (MI)**

Publicly available scRNA-seq datasets were obtained from the Gene Expression Omnibus (GEO). We analysed dataset GSE163956 (mouse model of MI)<sup>1</sup>. Data processing and analysis were performed using the Scanpy library (v.1.11) in Python. A standardised workflow was applied across datasets. After quality control and filtering based on gene counts and mitochondrial content, the data were normalised, log-transformed, and reduced to highly variable genes selected setting flavor to `seurat_v3`. Dimensionality reduction was then performed using PCA with 30 components, followed by UMAP projection and Leiden clustering to identify cell populations, which were annotated using canonical marker genes. Finally, the expression of MAP1S was analysed across cell types and biological conditions.

#### **Isolation and culture of neonatal rat cardiomyocytes**

Neonatal rat cardiomyocytes (NRCMs) were isolated from 2-3-day-old Sprague-Dawley rat neonates' hearts using enzymatic dissociation as previously described by our laboratory<sup>2</sup>. Briefly, the ventricles were digested in ADS buffer (116 mM NaCl, 20 mM HEPES, 1 mM NaH<sub>2</sub>PO<sub>4</sub>, 5.5 mM glucose, 5.5 mM KCl, 1 mM MgSO<sub>4</sub>; pH 7.35) containing 0.6 mg/ml collagenase A (Roche) and 0.6 mg/ml pancreatin (Sigma) at 37°C in a shaking water bath. Cardiac fibroblasts were separated by plating cells on 10-mm tissue-culture dishes and incubated in a humidified incubator at 37°C with 5% CO<sub>2</sub> for 1 hour. This process allows the fibroblasts to attach to the culture dish and to leave non-adherent cell, predominantly cardiomyocytes, in the media. Cells were plated on BD Falcon Primaria plates in medium containing 68% DMEM, 17% M199, 10% horse serum, 5% FBS, 2.5 µg/ml amphotericin B and 1 µM 5-bromo-2-deoxyuridine (BrdU). After 24-hours incubation at 37°C, cells were washed twice with PBS and maintained in a maintenance medium containing 80% DMEM, 20% M199, 1% FBS, 2.5 µg/ml amphotericin B, and 1 µM BrdU before being used in experiments.

#### **MAP1S overexpression and gene silencing in NRCMs**

Adenoviral vector was used to overexpress human MAP1S cDNA. Adenovirus expressing β-galactosidase (AdLacZ) was used for control. Human MAP1S cDNA was obtained from MyBioSource (plasmid #MBS1267628). The construct was cloned to an adenovirus vector pAd-CMV-DEST (Invitrogen) using a Gateway cloning system. NRCMs were transduced with adenovirus at a multiplicity of infection (MOI) 25 in maintenance media.

The siRNA-mediated gene silencing approach was used to knockdown MAP1S in NRCMs. siRNA targeting rat MAP1S was obtained from Sigma (Sigma, #SASI Rn02-00215332). Control siRNA was also obtained from Sigma (Sigma, SIC001). MAP1S or control siRNA were transfected to NRCMs using Dharmafect #1 transfection reagent (Dharmacon) following the protocol recommended by the manufacturer.

#### **Western blots**

Protein was extracted from the heart tissues or NRCMs using RIPA buffer supplemented with proteinase inhibitors. Protein concentration was measured using a Pierce BCA protein assay kit (Thermo Fisher Scientific) as per the manufacturer's instructions. An equal amount of protein sample (30 µg/sample) was separated by 8-15% SDS-PAGE, transferred to PVDF membranes, and blocked in 1-5% BSA or non-fat milk. Membranes were incubated respectively with primary and secondary antibodies. The blot bands were developed by ECL western blotting detection reagent (GE Healthcare) and imaged using ChemiDoc XRS+ imaging system (Bio-Rad). A list of antibodies used is available in Supplementary Table 3.

#### **Analysis of cardiomyocyte apoptosis using TUNEL assay**

TUNEL assay was used to determine cardiomyocyte apoptosis. NRCMs overexpressing or silencing MAP1S were treated with H<sub>2</sub>O<sub>2</sub> (200 µM, 2 hours) to induce oxidative stress and apoptosis. We used the In situ Cell Death Detection Kit (Roche) to detect TUNEL positive cells according to the manufacturer's recommended protocol. In each experimental group, 20 images were randomly captured. Apoptotic nuclei were quantified as percentage of TUNEL-positive nuclei over total nuclei in each experimental group.

#### **GFP-LC3 puncta formation assay**

We used adenovirus expressing GFP-LC3 to monitor autophagosome formation. The plasmid containing GFP-LC3 construct was a gift from Tamotsu Yoshimori (Addgene plasmid # 21073 ; http://n2t.net/addgene:21073 ; RRID:Addgene\_21073)<sup>3</sup>. This construct was cloned to adenovirus vector pAd-CMV-DEST using a Gateway cloning system (Thermo Fisher Scientific). Following MAP1S overexpression or gene silencing, cells were treated with AdGFP-LC3. The following day, cells were treated with rapamycin (5 µM, Sigma) to induce autophagy and chloroquine (3 µM, Sigma) to inhibit lysosome activity. After two washings with PBS, cells were fixed with 4% paraformaldehyde for 10 minutes and counterstained with DAPI. Fluorescence images (20 images/group) were randomly acquired using a Zeiss fluorescence microscope (Carl Zeiss, Jena, Germany). Determination of autophagosome number was performed by manually counting GFP-LC3 puncta within each cardiomyocyte using ImageJ software.

#### **Tandem mCherry-GFP-LC3 assay**

Tandem fluorescent-tagged LC3 reporter (mCherry-GFP-LC3), was used to examine autophagic flux. The plasmid containing mCherry-GFP-LC3 was a gift from Anne Brunet (Addgene plasmid # 110060 ; http://n2t.net/addgene:110060 ; RRID:Addgene 110060)<sup>4</sup> and was cloned to pAd-CMV-DEST using a Gateway cloning system (Thermo Fisher Scientific). This reporter allows distinction between autophagosomes and autolysosomes due to the pH sensitivity of GFP. The GFP signal is quenched when autophagosomes fuse to lysosomes to form autolysosomes, while the mCherry signal remains stable. Following overexpression of MAP1S, cells were transduced with AdmCherry-GFP-LC3 for 24 hours. Cells were then treated with rapamycin (5 μM, Sigma) and chloroquine (3 μM, Sigma) for 2 hours, washed, fixed with 4% paraformaldehyde, and counterstained with DAPI. Fluorescence images (20 images/group) were randomly acquired using a Zeiss fluorescence microscope (Carl Zeiss, Jena, Germany). Quantification of puncta was performed manually using ImageJ. Autophagosomes were identified as GFP<sup>+</sup>/mCherry<sup>+</sup> (yellow) puncta, while autolysosomes were defined as GFP<sup>-</sup>/mCherry<sup>+</sup> (red) puncta. Autophagic flux was determined by calculating the ratio of red to yellow puncta per cardiomyocyte.

#### **LC3-II turnover assay**

LC3-II turnover was assessed by Western blot. To estimate the amount of LC3-II degraded in lysosomes, LC3-II levels (normalized to GAPDH) in untreated cells were subtracted from those in cells treated with rapamycin and chloroquine.

#### **YAP nuclear translocation**

YAP subcellular localisation was assessed using a GFP-YAP construct. The method used to generate the GFP-YAP adenovirus was previously described<sup>2</sup>. Following MAP1S silencing or overexpression, NRCMs were transduced with an adenovirus expressing GFP-YAP construct for further 24 hours. Following overnight incubation, NRCMs were washed with PBS, fixed with 4% paraformaldehyde, and then counterstained with DAPI. Images were acquired using a Zeiss fluorescence microscope (Carl Zeiss, Jena, Germany). GFP-YAP nuclear translocation was detected by identifying nuclei that co-localised with the green signal.

#### **YAP luciferase assay**

YAP activity was assessed by using the YAP-luciferase reporter assay. The construction of the Ad-GAL4-TEAD and Ad-UAS-Luc is described in our previous publication<sup>2</sup>. Following overexpression or silencing of MAP1S, NRCMs were transduced with adenoviruses expressing GAL4-TEAD and UAS-luciferase for further 24 hours. The next day, the luciferase signal was detected using luciferase detection reagent (Promega) following the manufacturer's instructions and measured with a Lumat LB9507 Tube Luminometer (Berthold).

#### **Mitochondria function analysis in vitro**

Real-time oxygen consumption rate (OCR) was analysed to determine mitochondrial function of cardiomyocytes lacking MAP1S. NRCMs at density of $3 \times 10^4$ cells in 80 μl maintenance media were seeded onto laminin coated 96-well plate and then treated with siMAP1S or siControl for 72 hours. The OCR was measured in untreated and H<sub>2</sub>O<sub>2</sub> treated (200 μM, 2 hours) NRCMs using Seahorse XF Cell Mito Stress Test Kit (Agilent), according to the manufacturer's manual. The OCR was normalised to protein concentration of each treatment group. Based on the measured OCR values, the basal OCR, oxygen consumption linked to ATP production, level of non-ATP-linked oxygen consumption (proton leak), maximal and spare respiration capacity, and nonmitochondrial oxygen consumption were obtained.

#### **Myocardial infarction (MI) model**

Myocardial infarction was induced by permanent ligation of the left anterior descending (LAD) coronary artery as previously described<sup>5</sup>. Twelve-weeks-old MAP1S<sup>-/-</sup> and wild-type C57BL/6J mice were used for MI studies. Mice were anaesthetised using 3% isoflurane and oxygen at a flow rate of 1l/min, followed by an intraperitoneal injection of buprenorphine (0.1 mg/kg body weight) to provide pre-operative analgesia. Mice were then intubated and mechanically ventilated at a rate of 200 breaths per minute and a tidal volume of 0.1 ml. During the surgery, mice were maintained in a stable plane of anaesthesia with 1.5-3% isoflurane in 100% oxygen. The heart was exposed through a left thoracotomy at the fourth intercostal space. The LAD was ligated above its bifurcation approximately 1 mm below the edge of the left atrial appendage, using an 8-0 suture (ETHILON). Successful LAD occlusion was confirmed by discoloration of the myocardium distal to the ligation site. To determine the effect of modRNA-mediated MAP1S overexpression after MI, 50 μg/heart of either MAP1S or control luciferase modRNA was injected directly into the anterior wall of the left ventricle immediately after LAD ligation. Sham-operated (sham) animals were subjected to the same surgical procedures except that the suture was not tied around the LAD coronary artery. The thoracotomy and skin were sutured closed in layers. Mice were allowed to recover at 30 °C and closely monitored for the following 6 hours, before being returned to normal housing.

#### **Cardiac troponin I assay.**

To confirm the presence and extent of MI, serum cardiac troponin I (cTnI) levels were measured 24 hours post-MI. Blood samples were collected from the lateral tail vein. Prior to sampling, the tail was anesthetized with a topical 2.5% lidocaine/prilocaine cream, and mice were placed in restraint tubes on a warming pad to facilitate vasodilation. Following skin disinfection with povidone-iodine solution, a small transverse incision was made over the vein, and 40 µl of blood was collected and transferred to a tube containing 40 µl of 3.2% sodium citrate to prevent coagulation. Samples were centrifuged at 8,000 rpm for 6 minutes to isolate the plasma. Plasma cTnI levels were measured using a high-sensitivity mouse cardiac troponin I ELISA kit (Life Diagnostics), following the manufacturer's instructions. The absorbance was read at 450 nm using a FLUOstar Omega multiplate reader (BMG LABTECH) and plotted on the standard curve to calculate the cTnI concentration of each sample (ng/mL).

#### **Echocardiography analysis**

Transthoracic echocardiography was performed to assess cardiac function, left ventricular wall thickness and chamber dimensions. Anaesthesia was induced by 2% isoflurane and then maintained at 1%. Images were acquired in parasternal long and short axis views using a Vevo 3100 Imaging System equipped with a 30 MHz linear transducer (MX400; FUJIFILM VisualSonics Inc., Canada). All acquired images were later analysed using Vevo LAB analysis software (FUJIFILM VisualSonics Inc., Canada). Using the parasternal long axis view, the LV trace tool was used to determine the LV ejection fraction. Measurements of the left ventricular internal diameter (LVID), posterior wall thickness (PW), and interventricular septum thickness (IVS) at end-systole and end-diastole were obtained using the leading-edge method over a period of 3 cardiac cycles. Relative wall thickness was calculated as $2 \times \text{PWd}/\text{LVIDd}$.

#### **Histology analysis**

At the end of MI experiments, mice were euthanized by cervical dislocation in accordance with the United Kingdom Animals (Scientific Procedures) Act 1986. Mouse hearts were excised, briefly washed in PBS, weighed, and fixed in 4% paraformaldehyde in PBS for 24 hours at 4°C. The tissues were processed overnight in a Leica ASP300 automated tissue processor. The hearts were embedded in paraffin wax and sectioned at 5 µm thickness into 6-8 levels starting from the apex, with 500 µm intervals between each level using an automated rotary Leica RM2255 microtome. Masson's trichrome staining was performed to assess infarct size. Histological sections from the 6-8 levels of the heart were stained using a standard protocol. Slides were imaged on a 3D Histech Panoramic 250 Flash II slide scanner and analysed with SlideViewer (v2.7) software. Infarct size in all of the MI models was calculated using a midline length measurement approach as described previously <sup>6</sup>. For both methods, infarcted regions were measured in all levels of histological sections spanning from the apex to the base. Myocardial infarct size was expressed as a percentage of the total left ventricle area or circumference, as appropriate for the respective method. To analyse cardiomyocyte size, haematoxylin and eosin staining was performed using a standard protocol. Cardiomyocyte cross-sectional area was measured in both the remote and border zones using SlideViewer (v2.7) software, selecting only cardiomyocytes that were sectioned transversely, as indicated by the presence of a centrally located nucleus. To assess apoptosis, heart sections were stained using TUNEL reagents (In situ Cell Death Detection Kit, Roche) according to the manufacturer's recommended protocol.

#### **Transmission electron microscopy**

Transmission electron microscopy was carried out to analyse the ultrastructure of the mouse heart tissues post-MI. Heart tissues were fixed in 2.5% glutaraldehyde and 0.1M HEPES buffer (pH 7.2) containing 4% formaldehyde. Tissues were then processed in 0.1 M cacodylate buffer (pH 7.2) with 1% osmium tetroxide and 1.5% potassium ferrocyanide for 1 hour before treatment with 0.1 M cacodylate buffer (pH 7.2) and 1% uranyl acetate for a further 1 hour. Heart tissues were then treated with 1% uranyl acetate for an additional hour. After the fixation and treatment, the tissues were dehydrated using ethanol and then embedded in TAAB 812 resin and polymerised at 60 °C for 24 hours. Ultrathin sections were cut using a Reichert Ultracut ultramicrotome and examined with an FEI Tecnai 12 Biotwin transmission electron microscope at 100 kV accelerating voltage. Images were acquired from randomly selected areas using a Gatan Orius SC1000 CCD camera. Sample processing and imaging were performed at the University of Manchester Bioimaging facility. Quantification autophagosomes and autolysosomes was performed using ImageJ software. Autophagosomes were identified based on their characteristic double-membrane structure with a visible intermembrane space, while autolysosomes were distinguished as single-membrane vesicles containing amorphous, electron-dense material <sup>7</sup>. Autophagic flux was determined by calculating the ratio of autolysosome to autophagosome per heart. Measurement of mitochondrial circumference and crista length was performed using ImageJ.

#### **Synthesis and transfection of modified RNA**

We generated templates for the synthesis of MAP1S and luciferase modified RNAs (modRNAs) by cloning human MAP1S cDNA and luciferase genes in plasmid containing T7 promoter, optimised 5' and 3' untranslated regions (UTRs), and a polyA tail. modRNAs were transcribed in vitro from these plasmid templates. In vitro modRNA synthesis was performed using previously described method (nucleotide composition 5 (ARCA10))<sup>8</sup>. The composition of the in vitro transcription reaction is shown in Supplementary Table 4. Reactions were carried out at 37°C for 6 hours. Following transcription, the reaction mixture was purified using MEGAclear kit (Thermo Fisher Scientific) as per manufacturer's instructions and treated with Antarctic phosphatase (New England BioLabs) to remove 5' triphosphates. The treated modRNA was then re-purified with the MEGAclear kit and precipitated by adding 1:10 volume of 5M ammonium acetate and 2.5 volumes of ice cold 100% ethanol and left at -20°C overnight. The following day, samples were centrifuged at 14,000 rpm for 15 minutes at 4°C. The RNA pellet was washed with 500 µl ice-cold 70% ethanol, air-dried, and resuspended in 20 µl nuclease-free water. modRNA concentration was determined using a NanoDrop 8000 spectrophotometer (Thermo Fisher Scientific).

For the in vivo studies, 50 µg of either MAP1S or luciferase modRNA was complexed with lipofectamine RNAiMAX transfection reagent (Life Technologies) according to manufacturer's recommendations, in a total volume of 30 µl. The transfection mixture was directly injected into the left ventricular myocardium using 0.3ml insulin syringe with a 30G needle. The needle was held in place for a few seconds to reduce the risk of the modRNA solution being ejected from the site of injection.

#### **In silico analysis**

To identify MAP1S-interacting partners, protein-protein interaction analysis was performed using publicly available databases including BioGRID<sup>9</sup>, HitPredict<sup>10</sup>, STRING<sup>11</sup>, IntAct<sup>12</sup>, Mentha<sup>13</sup>, WiKi-Pi<sup>14</sup>, and NCBI database (https://www.ncbi.nlm.nih.gov/). Non-human and non-experimentally validated interactors of MAP1S collected from these databases were excluded from further analysis. Next, expression analysis was performed in the Expression

Atlas database<sup>15</sup> to exclude MAP1S interactors not expressed in the human heart. Only proteins whose association with MAP1S have been confirmed by at least two pieces of experimental evidence were included in pathway analysis. Kyoto Encyclopedia of Genes and Genomes (KEGG)<sup>16</sup> was used to identify the clusters of genes of MAP1S interactors with their associated signalling pathways. In addition, to analyse the enriched pathways associated with the MAP1S interacting proteins, overrepresentation analysis was carried out in Reactome database<sup>17</sup>. A p-value < 0.05 was considered as significant.
